# The microbiota extends the reproductive lifespan by safeguarding the ovarian reserve

**DOI:** 10.1101/2024.09.13.612929

**Authors:** Sarah K. Munyoki, Julie P. Goff, Amanda Reshke, Erin Wilderoter, Nyasha Mafarachisi, Antonija Kolobaric, Yi Sheng, Steven J. Mullett, Gabrielle E. King, Jacob D. DeSchepper, Richard J. Bookser, Carlos A. Castro, Stacy L. Gelhaus, Mayara Grizotte-Lake, Kathleen E. Morrison, Anthony J. Zeleznik, Timothy W. Hand, Miguel A. Brieño-Enriquez, Eldin Jašarević

## Abstract

Infertility is a devastating condition affecting one in six people globally. In many cases, the underlying causes are unknown. Emerging evidence suggests that the microbiota influences reproduction, yet the mechanistic link between the microbiota, ovarian function, and length of the fertile lifespan remain unexplored. Here we report that the microbiota controls the length of the reproductive lifespan by maintaining the primordial follicle pool, a process mediated by microbiota-derived short chain fatty acids modulating gene regulatory networks crucial for the survival of the ovarian reserve. Dietary perturbation of the microbiota during a critical developmental window is sufficient to diminish the ovarian reserve, reduce oocyte retrieval, and impair preimplantation embryo viability, mirroring challenges in human fertility treatments. Targeted interventions to restore microbiota improve assisted reproductive outcomes, particularly when implemented early. These findings reveal a novel contribution of host-microbe interactions in mammalian reproduction and demonstrate that the microbiota impacts ovarian function and fertility.

## Introduction

Infertility is a devastating condition affecting one in six people globally, with its prevalence rapidly increasing worldwide (*1*). Similar alarming trends are reported across the animal kingdom, underscoring an urgent need to identify unifying principles and mechanisms governing reproductive health. Recent years have seen the emergence of the microbiota as a critical regulator of host physiology, including metabolism and immunity, and more recently, reproduction and pregnancy outcomes (*2–12*). The presence of specific bacteria is necessary for the sexual maturation and production of viable oocytes in invertebrate species (*13–17*). In mammals, antibiotic-mediated microbiota perturbations negatively impact male fertility (*18*). However, the contribution of the microbiota in female reproduction remains unexplored. This knowledge gap is particularly significant given growing recognition that metabolic disorders, such as obesity, are associated with both microbiota perturbations and reproductive dysfunction (*19–21*). We therefore examined the hypothesis that the presence of the microbiota controls ovarian function and reproductive fitness throughout the lifespan. Our study addresses a fundamental and longstanding question in reproductive biology regarding the preservation of the ovarian reserve, a finite and non-renewable pool of oocytes, providing insights into the contribution of the microbiota to female reproductive health.

## Results

### Microbiota controls the reproductive lifespan by maintaining the ovarian reserve

In the 1970s, researchers found that germ-free mice produced smaller litters than conventional mice (*22, 23*). Despite the potential to deepen our understanding of novel mechanisms regulating fertility, this link between the microbiota and mammalian reproduction remained unexplored for decades. Building on this early insight, we began by collating and analyzing lifetime breeding records from 493 litters of conventionally raised, murine-pathogen-free (MPF), and germ-free (GF) C57Bl/6N Tac mice. MPF mice produced a mean of 4.7 litters per lifespan, while GF mice produced a mean of only 2.5 litters (Fig. 1A). The average number of pups per litter was lower in GF mice (Fig. 1B). MPF mice continued to birth offspring up to the ninth litter. In contrast, GF mice became infertile by the fifth litter (Fig. 1, C and D). We performed undersampling with 1000 permutations to account for differences in sample numbers and confirm the robustness of these results (fig. S1). These data suggest that maintaining consistent litter sizes across multiple pregnancies, total lifetime number of offspring, and overall reproductive lifespan requires the microbiota in mice.

**Figure 1:**
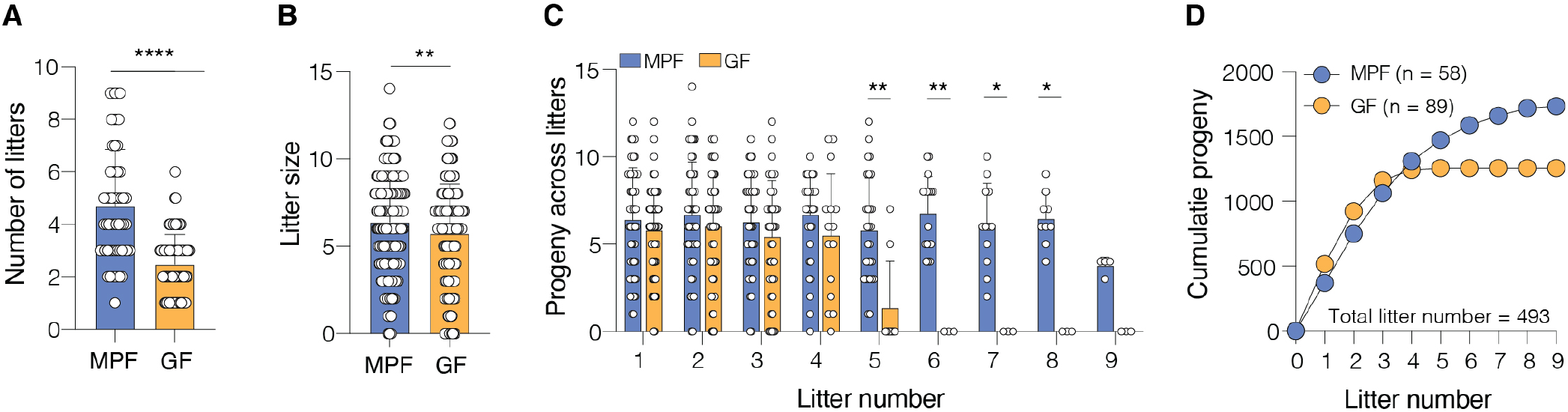
The microbiota extends the reproductive lifespan in mice. **(A)** Lifetime litter production in MPF (n = 58) and GF (n = 89) female mice. **(B)** Average litter size of MPF (n = 273) and GF (n = 220) litters. **(C)** Progeny per litter across sequential births. **(D)** Cumulative progeny over sequential litters in MPF and GF mice (total n = 493 litters). Data in (A-C) show mean ± SD. (A-B) analyzed by two-tailed Student’s t-test. (C) analyzed by mixed-effects ANOVA followed by Šidák’s test. *p < 0.05, **p < 0.01, ****p < 0.0001. MPF, murine pathogen free; GF, germ-free. See also fig. S1.

To understand the biological basis underlying effects of the microbiota on reproductive output, we examined reproductive tissues from 10-week-old adult (P70) MPF and GF mice (Fig. 2A). Histopathology of testes from MPF and GF male mice displayed normal spermatogenesis, including the presence of spermatids in the lumen of seminiferous tubules (fig. S2). In contrast, ovaries from GF female mice exhibited distinct abnormalities, including hemorrhagic lesions and fibrosis (Fig. 2B). The mammalian ovary is a dynamic organ characterized by follicles progressing through distinct developmental stages (*24*). All ovarian follicles are recruited from a non-renewable pool of primordial follicles, termed the ovarian reserve. This reserve is established in early life and represents the total reproductive potential in female mammals. Once activated, primordial follicles develop into primary, secondary, and preovulatory antral follicles, ultimately leading to ovulation and corpus luteum formation to support pregnancy or follicle cell death, termed atresia (*25– 31*). Quantification of follicle stages revealed a specific effect of the microbiota on the ovarian reserve (Fig. 2C) (*32, 33*). Primordial follicles in GF ovaries were reduced by 50% compared to MPF females (MPF: 766 vs. GF: 376) (Fig. 2D).

**Figure 2:**
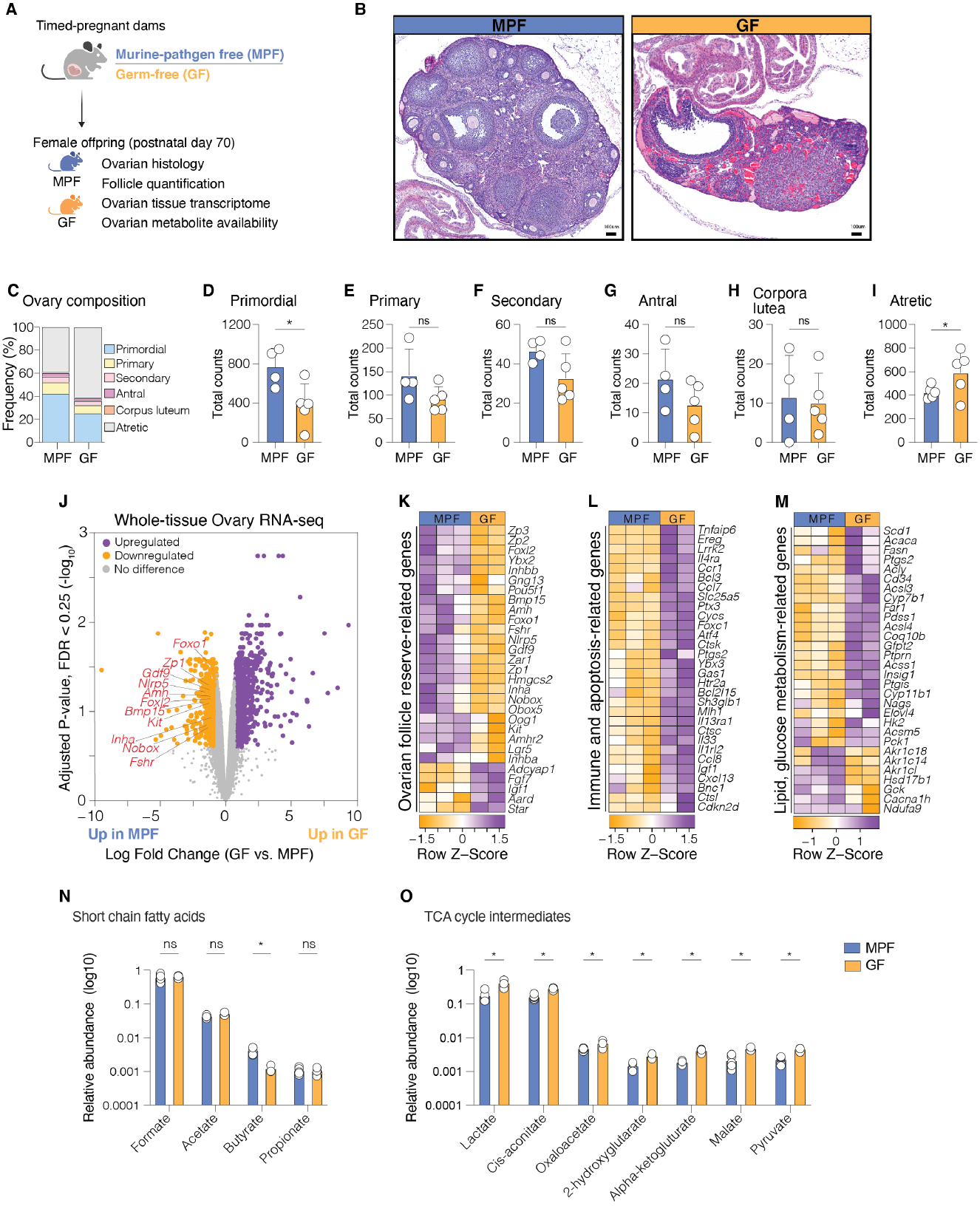
Gut microbiota impact reproductive capacity through regulation of ovarian follicle dynamics. **(A)** Experimental design: Ovarian analysis of 10-week-old female offspring from MPF and GF dams. **(B)** Representative ovarian histology (H&E staining, scale bars: 100 μM). **(C-I)** Ovarian follicle quantification: (C) Relative ovarian follicle composition, (D) primordial, (E) primary, (F) secondary, (G) antral follicles, (H) corpora lutea, and (I) atretic follicles. **(J)** Volcano plot of differentially expressed ovarian genes (GF vs. MPF). Purple: upregulated in GF; orange: downregulated. Key ovarian reserve genes annotated. Adjusted p-value, FDR < 0.25. **(K-M)** Heatmaps of gene expression related to (K) ovarian reserve, (L) immunity, and (M) metabolism. (**N**) Short-chain fatty acids in ovaries. (**O**) TCA cycle intermediates in ovaries. n = 3-5/group (C-H, N, O). Data: mean ± SD. Statistics: two-tailed Student’s t-test with Tukey’s test (C-G), Poisson regression with Tukey’s test (H-I). Data are representative of two independent experiments. Each point represents an individual non-littermate female mouse. Groups: MPF, murine pathogen free; GF, germ-free. *p < 0.05, **p < 0.01, ****p < 0.0001. See also fig. S2 to S4.

Primary, secondary, and antral follicle counts were comparable between groups (Fig. 2, E to G), suggesting that the gonadotropin-dependent phase of follicle development is intact in GF mice (*34, 35*). The presence of corpora lutea was similar in both groups, indicating that ovulation is also intact (Fig. 2H). However, GF ovaries showed an increased number of atretic follicles compared with MPF females, suggesting that the reduced primordial follicle pool is associated with increased follicle atresia in the absence of microbiota (Fig. 2I).

We performed whole-tissue RNA sequencing (RNAseq) of P70 MPF and GF ovaries to elucidate the molecular mechanisms underlying the observed ovarian phenotype. This analysis identified over 600 differentially expressed genes, including downregulation of genes encoding factors involved in primordial follicle quiescence, activation, and survival (e.g., *Kit, Amh, Nobox, Gdf9*) (Fig. 2J) (*36–40*). Gene ontology analysis of differentially expressed genes showed an overrepresentation of pathways involved in the ovarian reserve, immunity, metabolism, and tissue remodeling (Fig. 2, K to M). To explore the functional significance of the observed upregulation of genes involved in tissue remodeling (*Col1a1, Col3a1, Vcan*), we performed Masson’s trichrome staining, showing extensive collagen fiber deposition in GF ovaries and absent in MPF ovaries (fig. S3, A to B) (*41*). These transcriptional changes were accompanied by metabolic changes, including decreased availability of microbial metabolites and increased abundance of all measured TCA cycle intermediates in GF compared with MPF ovaries (Fig. 2, N to O).

Reduced reproductive capacity affects the entire body, including the neural circuits related to reproduction (*42*). To assess whether alterations in the hypothalamus contribute to the ovarian phenotype observed in GF mice, we measured the expression of *Kiss1* and *Pdyn* in the arcuate nucleus of the hypothalamus in P70 MPF and GF females. These genes are co-expressed in KNDy (Kisspeptin/Neurokinin B/Dynorphin) neurons, which regulate fertility and pregnancy (*42–45*). Using quantitative PCR (qPCR), we found no differences in *Kiss1* or *Pdyn* mRNA transcript levels between MPF and GF female mice (fig. S4, A to B). This finding, combined with our characterization of ovarian phenotypes (Fig. 2), suggests that the reproductive deficits in GF mice stem from effects on ovarian tissue rather than alterations to neural circuits regulating reproduction.

### Postnatal microbiota maturation shape ovarian follicle dynamics across development

To understand the developmental origins of the adult ovarian phenotype, we quantified follicle stages across time points that capture critical phases of ovary postnatal maturation (P7, 21, 28, 70) (Fig. 3A) (*46*). Based on our characterization of the adult ovarian phenotype, we reasoned that GF mice start with a lower reserve early in life, which is then depleted more rapidly across repeated cycles of ovulation.

**Fig. 3.**
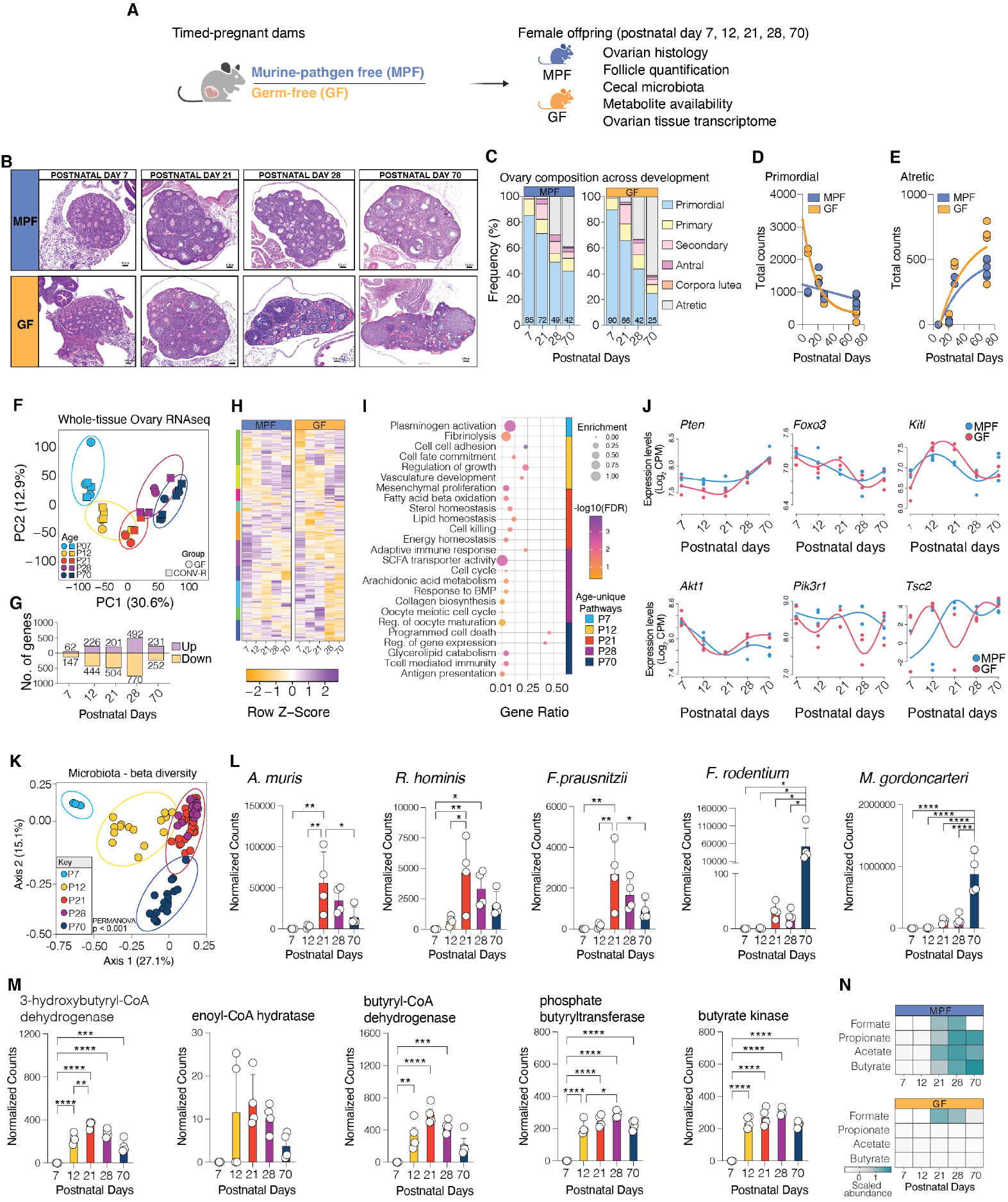
Postnatal gut microbiota preserves the ovarian reserve through developmental gene regulation. **(A)** Experimental design: Parallel ovarian and microbiota development in MPF and GF mice (postnatal days 7-70). **(B)** Ovarian histology at 7, 21, 28, and 70 days (H&E, scale: 100 μM). **(C-E)** Follicle dynamics in MPF vs. GF: (C) Composition across development, (D) primordial, (E) atretic. **(F)** PCA of ovarian transcriptomes. **(G)** Differential gene expression in GF vs. MPF ovaries (FDR < 0.25). **(H)** Heatmap of differentially expressed genes. **(I)** Bubble plot of enriched GO terms for age-specific genes downregulated in GF, highlighting microbiota-dependent impacts on metabolites, immune, and cell cycle programs at specific postnatal timepoints. **(J)** Time series reconstruction of PI3K/AKT/mTOR pathway gene dynamics. **(K)** PCoA of cecal microbiota (16S rRNA) in MPF mice. **(L)** SCFA-producing bacterial species abundance (shotgun). **(M)** Microbial SCFA production gene counts (shotgun). **(N)** Cecal SCFA abundance heatmap. n = 3-5 per group/timepoint (A-J, N); n = 3-4 MPF/timepoint (K-M), with 1 animal used per litter for analysis. Data: mean ± SD. Statistics: (C) nonlinear regression, (L,M) mixed-effects ANOVA with Bonferroni’s test, (K) PERMANOVA (1000 permutations). Data are representative of three independent experiments. Each point represents an individual non-littermate female mouse. Groups: MPF, murine pathogen free; GF, germ-free; SCFA, short-chain fatty acid. *p < 0.05, **p < 0.01, ****p < 0.0001. See figs. S5 to S8.

Such a scenario could account for differences in ovarian reserve between MPF and GF mice. Testing this, our analysis revealed significant postnatal effects of the microbiota on ovarian follicle dynamics (Fig. 3, B and C). GF mice had more primordial follicles at P7 (MPF: ∼1400 vs. GF: ∼2100) (Fig. 3D). However, this early advantage was rapidly lost. By P21, the number of primordial follicles in GF mice started to decline (MPF: ∼1300 vs. GF: ∼900) and continued through P70 (MPF: ∼700 vs. GF: ∼400) (fig. S5A). Regression analysis showed that the rate of primordial follicle loss in GF mice followed an exponential decay pattern, in contrast to the linear decline in MPF mice (Fig. 3D). Consistent with a higher rate of primordial follicle activation, we observed a transient increase in primary follicles in GF mice between P21 and P28 (fig. S5). However, this increased recruitment did not correlate with higher secondary and antral follicle counts at subsequent time points. Instead, we observed an increase in atretic follicles in GF mice (Fig. 3E and fig. S5), suggesting that primordial follicles prematurely activate, fail to mature, and undergo atresia in the absence of microbiota.

We next investigated the molecular consequences to an ovary developing without the microbiota. Whole-tissue RNAseq analysis of ovaries at each time point revealed distinct clustering patterns based on microbiota status and age, with the impact of microbiota becoming more pronounced at later time points (Fig. 3F). Principal component analysis showed a clear separation of samples by age along PC1 (30.6% of variation) and by microbiota status along PC2 (12.9% of variation). Differences in gene expression progressively increased from P7 (∼200 genes) to P28 (∼1300 genes), indicating that the effect of the microbiota on ovarian transcription becomes more pronounced as the female reproductive system matures (Fig. 3, G and H). Gene ontology analysis revealed that the absence of microbiota affects distinct biological processes at each developmental time point.

Downregulated genes in GF mice (and upregulated in MPF mice) were associated with cell fate commitment, oocyte maturation, SCFA transport, immunity, and energy homeostasis (Fig. 3I), suggesting microbiota dynamically influence key transcriptional programs in ovarian tissue throughout postnatal development.

Further analysis of genes critical for regulating primordial follicle dynamics revealed several key players influenced by microbiota (table S1 and fig. S6) (*27*). Of particular interest were time-series expression patterns of genes involved in the PI3K/Akt/mTOR signaling pathways, critical for regulating primordial follicle activation, quiescence, and survival (Fig. 3J) (*47–49*). *Pten, Foxo3*, and *Tsc2*, involved in suppressing follicle activation, and *Akt1, Kit/Kitl*, and *Pik3r1*, which promote activation and oocyte survival (*36, 39, 47, 49, 50*), showed altered temporal expression patterns in GF mice from P7 to P70. These findings suggest that the microbiota influence ovarian follicle dynamics and gene expression patterns throughout postnatal development, thereby supporting the establishment and preservation of the ovarian reserve.

To understand how these ovarian changes associate with the developing microbiota, we performed 16S rRNA gene profiling of cecal contents of MPF mice from P7 to 70. Principal coordinates analysis revealed a clear separation of samples by age, with P7 and P70 at opposite extremes (Fig. 3K). The early phases of the microbiota weaning transition began at P12, characterized by increased microbial diversity and a shift from *Lactobacillaceae* dominance to obligate anaerobes like *Clostridia* and *Bacteroidaceae* (*51*) (Fig. 3K and fig. S7). This transition coincides with the onset of solid food consumption, which introduces dietary polysaccharides and complex carbohydrates to the intestinal tract, enabling bacterial fermentation of these substrates into short-chain fatty acids (SCFAs) (*52*). Metagenomic shotgun sequencing confirmed an expansion of bacterial species and microbial metabolic pathways involved in SCFA production, starting at P12 and continuing into adulthood (Fig. 3, L and M). Targeted metabolomics confirmed increased cecal SCFA availability starting at P12 and persisting into adulthood in MPF but not GF mice (Fig. 3N and fig. S8, E to L). These differences were independent of body growth trajectories in MPF and GF mice (fig. S8, A to D). These results reveal that the postnatal maturation of the microbiota, particularly the increase in SCFA-producing bacteria during the microbiota weaning transition, plays a crucial role in regulating ovarian follicle dynamics, including maintenance of the primordial follicle pool.

### Postnatal microbiota colonization prevents germ-free ovarian reserve depletion

Having observed an association between postnatal maturation of the microbiota and ovarian follicle dynamics, we next assessed the requirement for the microbiota at specific postnatal periods in this process. We introduced microbiota from MPF donors to GF mice at two critical time points: the day of birth (P0) and the onset of the microbiota weaning transition observed in our MPF mice (P12) (Fig. 4A). We confirmed successful engraftment in colonized dams by 16S rRNA marker gene profiling and targeted metabolomics, showing recovery of microbial diversity, restoration of SCFA-producing *Clostridia*, and SCFAs levels comparable to MPF dams (fig. S9). Quantification of all follicle stages in P70 ovaries revealed that both P0 and P12 colonization rescued primordial follicle counts to MPF counts (Fig. 4, B to E). P0 and P12 colonization reversed morphological phenotypes observed in GF mice, restoring collagen deposition and fibrosis patterns to match those of MPF ovaries (fig. S10E). This rescue effect was specific to the primordial follicle pool, as growing follicles and corpora lutea numbers remained consistent across all groups (fig. S10, A to D). Using qPCR, we confirmed that colonization at either P0 or P12 restored the expression of ovarian reserve marker genes (*Kit, Amh, Nobox, Gdf9*) to levels similar to MPF mice (Fig. 4, F and G, fig. S10F) (*36, 37, 39, 40, 53–55*). Beyond local effects on ovaries, colonization at P0 or P12 normalized the cecum-to-body mass ratio, microbiota composition, and SCFA levels, aligning them with those of MPF females (Fig. 4H and fig. S11). Notably, the similarity in outcomes between P0 and P12 colonization suggest that presence of the microbiota as late as P12 is sufficient to prevent the ovarian reserve depletion observed in GF adult mice.

**Fig. 4.**
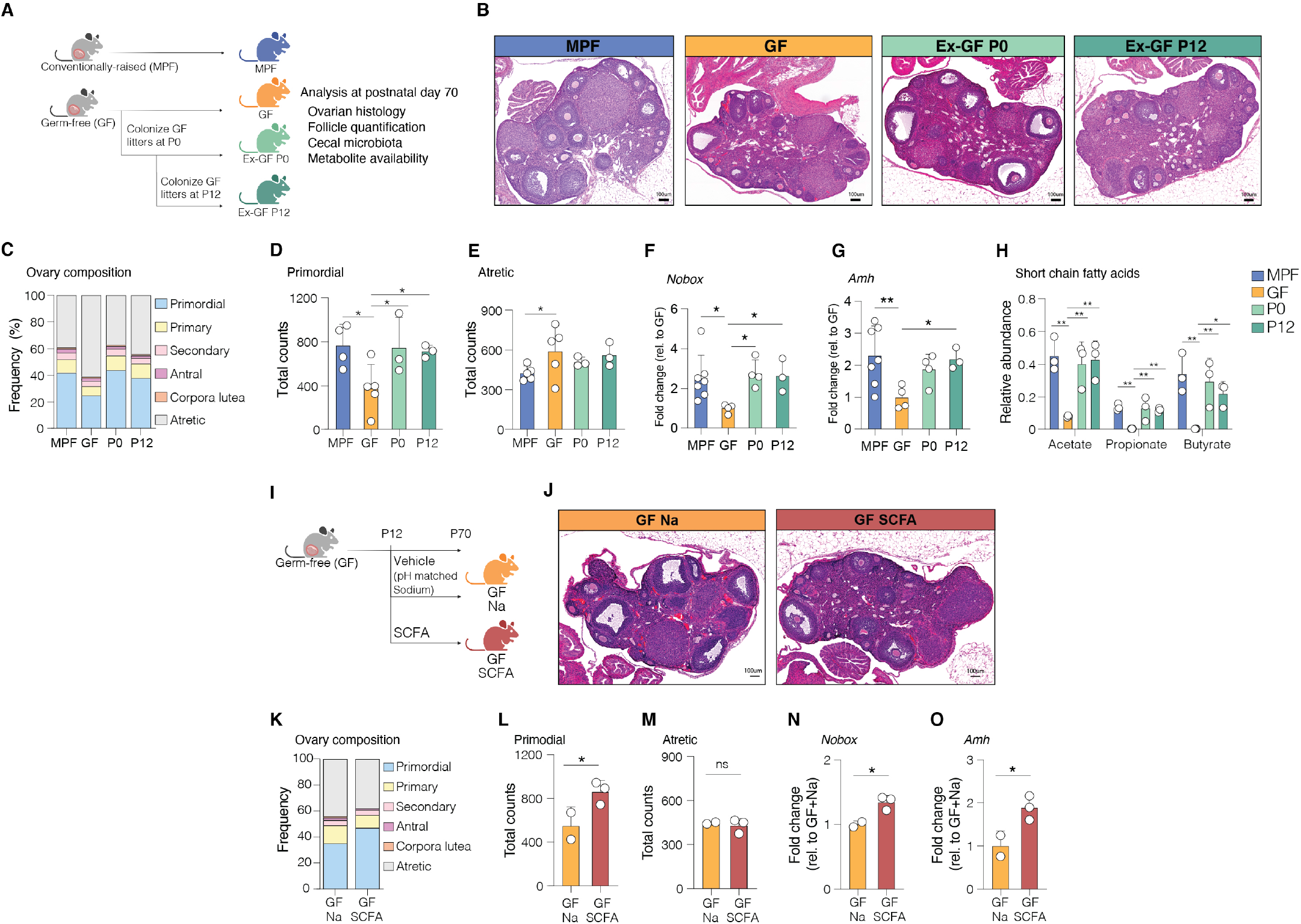
Microbial-derived metabolites mediate the preservation of the ovarian reserve. **(A)** Experimental design: GF mice colonized at birth (P0) or weaning onset (P12, reflecting the earliest stages in bloom of obligatory anaerobes and SCFA-producers). Analysis was conducted at postnatal day 70. **(B)** Ovarian histology: MPF, GF, P0-colonized, P12-colonized mice (H&E, scale: 100 μM**). (C-E)** Follicle quantification: (C) Ovary composition, (D) primordial, (E) atretic follicles. **(F, G)** qPCR of Nobox and Amh mRNA transcript levels. **(H)** Cecal SCFA abundance. **(I)** Experimental design: GF mice given SCFA or Vehicle sodium-matched water (P12-70). **(J)** Ovarian histology: GF+Na, GF+SCFA (H&E, scale: 100 μM). **(K-M)** Follicle quantification post-reconstitution: (J) Ovary composition, (K) primordial, (L) atretic follicles. (M) qPCR of Nobox and Amh mRNA transcript levels in GF+Na and GF+SCFA mice. n = 2-5 mice/group, with 1 animal used per litter for analysis. Data: mean ± SD. Statistics: one-way ANOVA with Fisher’s LSD (D, F and G). Poisson regression with Tukey’s test (E). Data are representative of three independent experiments. Each point represents an individual non-littermate female mouse. Groups: MPF, murine pathogen free; GF, germ-free; P0, colonized at birth; P12, colonized at weaning onset; SCFA, short-chain fatty acids. *p < 0.05, **p < 0.01. See also figs. S9 to 12.

### Short chain fatty acids mediate host transcription in ovarian reserve maintenance

Our results suggest that factors secreted by the microbiota as exerting an effect on ovarian follicle dynamics. We hypothesized that SCFAs are one class of microbial metabolites involved in the maintenance of the ovarian reserve. Starting at P12, mirroring the postnatal timepoint of SCFA availability, GF females received either an SCFA mix (67.5 mM acetate, 40 mM butyrate, 25.9 mM propionate) or pH and sodium-matched solution in their drinking water (Fig. 4H). Histological analysis of ovaries at P70 showed increased presence hemorrhagic lesions in ovaries in vehicle-supplemented GF females (GF Na) compared with SCFA-supplemented GF females (GF SCFA) (Fig. 4I). Quantification of follicles revealed an increase in the proportion of primordial follicles in SCFA-supplemented GF ovaries, while the proportions of other follicles were similar between groups (Fig. 4J, fig. S12, A to D). Further analysis showed a significant increase in primordial follicle counts in GF SCFA ovaries compared to GF Na ovaries, with no difference in atretic follicle counts (Fig. 4, K to M). Using qPCR, we found that SCFA supplementation in GF ovaries significantly increased the expression of *Nobox* and *Amh* compared to vehicle-treated GF ovaries (Fig. 4, N and O, fig. S12, E and F). *Nobox* encodes a transcription factor critical for early folliculogenesis and primordial follicle maintenance, while *Amh* inhibits premature primordial follicle activation (*36, 53*). These findings suggest that the microbiota may influence the expression of genes involved in ovarian follicle maintenance, in part through the production of SCFAs.

### Diet-induced microbiota disruption impairs the ovarian reserve and fertility outcomes

Western-style diets are increasingly associated with rising rates of infertility (*56–58*). These diets induce perturbations to the microbiota, which in turn increase the risk for poor health outcomes, including reproductive disorders (*21, 59, 60*). We previously demonstrated that high-fat, low-fiber diets rapidly alter gut microbiota composition, decreasing SCFA-producing bacteria and SCFA availability. These chronic changes impact maternal physiology, reproductive outcomes, and offspring development (*61–63*), and are associated with systemic inflammation and metabolic dysfunction that negatively impact reproductive function. Importantly, high-fat diets impair ovarian function in mice even in the absence of obesity, suggesting that obesity alone does not fully explain ovarian dysfunction (*58, 64–67*).

To further investigate the role of the microbiota in ovarian function, we examined whether diet-induced microbiota disruption was sufficient to replicate the ovarian phenotype observed in GF mice. We employed our established high-fat, low-fiber (Hf-Lf) diet model, known to alter gut microbiota composition and function (*61–63*). MPF female mice were fed either a control diet (CON) or Hf-Lf diet from P21 to P70 (Fig. 5A). We chose this timeframe to specifically disrupt the microbiota weaning transition and assess its effects on ovarian follicle dynamics in adulthood.

**Figure 5:**
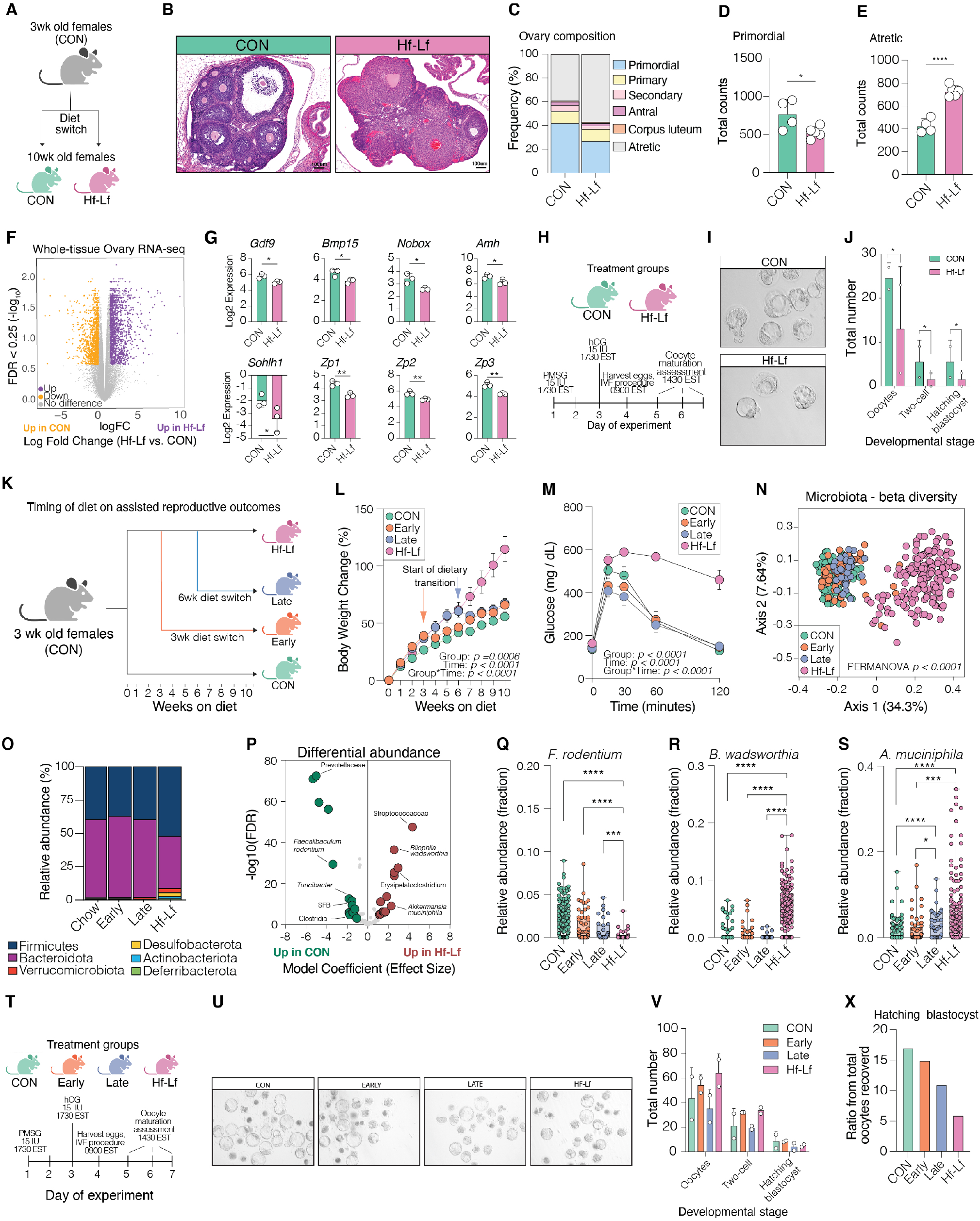
Diet-induced microbiota perturbation impairs ovarian reserve and fertility. **(A)** Experimental design: 3-week-old females fed control (CON) or high-fat, low-fiber (Hf-Lf) diet until 10 weeks. **(B-E)** Ovarian analysis at 10 weeks: (B) Histology (H&E, scale: 100 μM), (C) Ovary follicle composition, (D) primordial, (E) atretic follicles. **(F)** Whole-ovary RNA-seq: Differential gene expression (Hf-Lf vs. CON). **(G)** Expression of key ovarian function genes. **(H-J)** Assisted reproduction outcomes: (H) Experimental timeline, (I) Embryo images, (J) Developmental stage quantification. **(K)** Experimental design timing of dietary intervention experiment. **(L)** Body weight changes. **(M)** Glucose tolerance test. **(N)** Gut microbiota beta diversity. **(O)** Bacterial phyla abundance. **(P)** Differential abundance of bacterial taxa (CON vs. Hf-Lf). **(Q-S)** Abundance of *F. rodentium, B. wadsworthia, A. muciniphila* across diet groups. **(T-X)** Diet intervention reproductive outcomes: (T) Timeline, (U) Embryo images, (V) Developmental stages, (X) Hatching blastocyst ratio. n = 3-5/group (B-G), n = 12-20/group (L-S), oocytes pooled from 5 females per group/replicate with 2-3 independent replicates (J,V,X). Data: mean ± SD. Statistics: (E and J) analyzed using Poisson regression followed by Tukey test. (F,G) analyzed with FDR <0.25. (L,M) analyzed using two-way ANOVA followed by Tukey’s test. (N) Beta diversity analyzed using PERMANOVA with 1000 permutations. (P-S) Differential abundance analyses conducted using MaAsLin2 (Microbiome Multivariable Association with Linear Models 2). (J,V) analyzed using Poisson distributions. Groups: CON, continuous control diet; Hf-Lf, continuous high-fat low-fiber diet; Early, early diet switch; Late, late diet switch. *p < 0.05, ****p < 0.0001. See also figs. S12 to 13.

Hf-Lf feeding reduced primordial follicle counts comparable to that observed in GF mice at P70 (Hf-Lf: 528 vs. GF: 376), along with a significant increase in atretic follicles (Fig. 5, B to E). The numbers of growing follicles and corpora lutea were comparable across all groups, indicating that Hf-Lf effects are specific to the primordial follicle pool at this timepoint (fig. S13). RNAseq analysis of P70 Hf-Lf and CON ovaries revealed disruption to gene expression patterns in Hf-Lf mice similar to those observed in GF mice, including genes involved in ovarian follicle activation, survival, and atresia (Fig. 5, F and G).

To assess the functional significance of diet-induced microbiota perturbations, we employed superovulation and *in vitro* fertilization (IVF) to evaluate the effects on female fertility and embryo development (Fig. 5H). Hf-Lf mice produced fewer oocytes upon ovarian stimulation, and following IVF, their embryos showed reduced developmental competence, mirroring common and long-standing challenges seen in human IVF treatments for obese patients (*68, 69*) (Figure 5, I and J).

Recent clinical studies report that lifestyle interventions in obese patients often improve metabolic outcomes without enhancing fertility (*70*). We hypothesized that this discrepancy might be due to interventions occurring after significant exhaustion of the ovarian reserve, limiting their clinical benefit. To test this in our model, we investigated whether the timing and duration of Hf-Lf-induced microbiota disruption critically influenced reproductive outcomes. We established four groups of P21 MPF females: continuous CON, continuous Hf-Lf, Early intervention (3 weeks Hf-Lf followed by 7 weeks CON), and Late intervention (6 weeks HfLf followed by 4 weeks CON) (Fig. 5K). Metabolic assessment revealed that continuous Hf-Lf females showed the most substantial weight gain and glucose intolerance, while both intervention groups exhibited recovery of body weight and glucose homeostasis (Fig. 5, L and M). Microbiota analysis showed distinct clustering based on diet, with intervention groups shifting towards the CON profile after dietary change (Fig. 5N). Continuous Hf-Lf feeding reduced *Firmicutes* while increasing *Bacteroidota, Verrucomicrobiota*, and *Desulfobacterota* abundance, with both intervention groups showing restoration of community proportions comparable to CON females (Fig. 5O). Differential abundance analysis revealed diet-specific changes in key bacterial species, including *F. rodentium, B. wadsworthia*, and *A. muciniphila*, with *A. muciniphila* showing partial recovery in the Late intervention group, emphasizing the impact of timing on microbiota composition (Fig. 5, Q to S, fig. S14).

To determine whether timing of dietary intervention influenced reproductive outcomes, we used superovulation and IVF to assess oocyte quality and embryo development (Fig. 5T). The Early intervention group showed greater improvements in oocyte yield and blastocyst formation rates than the Late intervention group, approaching levels seen in CON mice (Fig. 5, U to X). These data support the conclusion that diet-induced microbiota disruption has lasting effects on ovarian follicle dynamics and fertility. Reproductive outcomes may be improved by dietary interventions aimed at correcting microbiota composition, with the timing of interventions critical; earlier interventions show more substantial benefits.

## Discussion

This study reveals a novel contribution of the microbiota on ovarian function and reproductive longevity in female mice. We demonstrate that the absence of microbiota leads to diminished reproductive output through disruption of metabolic and transcriptional programs, resulting in accelerated depletion of the primordial follicle pool that constitutes the ovarian reserve. Our findings highlight the critical role of microbial colonization timing and diet-induced microbiota perturbations in modulating ovarian function, fertility outcomes, and reproductive capacity. From an evolutionary perspective, our results raise fundamental questions about the reliance of the microbiota on mammalian reproductive processes. This dependence may point to host-microbiota co-evolution that shaped reproductive strategies across species. These results also suggest that the microbiota may serve as a barometer of environmental conditions, in which reproduction is optimized when conditions favor offspring survival and fitness.

Central to our findings is the identification of microbial metabolites, such as SCFAs, in modulating ovarian follicle dynamics. We propose microbiota-derived metabolites may impact the expression of genes involved in regulating the quiescence, activation, survival, and atresia of the primordial follicle pool. This hypothesis is supported by our observations that the absence of microbial signals leads to dysregulation of key pathways, resulting in premature follicle activation and subsequent atresia. Most notably, SCFA supplementation in germ-free mice was sufficient to influence the primordial follicle pool and the expression of transcripts involved in promoting primordial follicle maintenance.

The specific impact of the microbiota on the primordial follicle pool is particularly intriguing. Primordial follicle position along the perimeter of ovarian cortex may expose them continuously to circulating signals influenced by the microbiota. Moreover, the dormant state could make the follicles especially vulnerable to alterations in energy metabolism or oxidative stress, both of which are influenced by the microbiota. Our colonization experiments revealed a critical postnatal period for microbiota-dependent regulation of ovarian function. The striking rescue of ovarian phenotypes by microbial colonization as late as postnatal day 12 coincides with the transition in microbial maturation, a period characterized by significant changes in microbial composition and metabolite production. We propose that this developmental stage is crucial for establishing a previously uncharacterized communication between the microbiota and the ovary, with lasting consequences for reproductive health.

By elucidating the contribution of the microbiota in safeguarding the ovarian reserve and extending the reproductive lifespan, we solve a long-standing biological mystery regarding the reproductive ability of germ-free mice. These findings open new possibilities for preserving and potentially extending female fertility through targeted modulation of microbial communities. Future research should explore the relationship between microbiota composition and ovarian reserve in individuals with reproductive disorders, including diminished ovarian reserve, premature ovarian failure, or polycystic ovarian syndrome. Additionally, investigating the efficacy of microbiota-directed interventions, such as probiotics or prebiotics, on fertility outcomes in clinical settings represents a promising avenue for developing novel therapeutic approaches. These insights not only advance our understanding of reproductive biology but may offer promising avenues for preserving female fertility.

## Acknowledgements

We acknowledge Drs. William MacDonald and Amanda Poholeck in the Children’s Hospital of Pittsburgh Health Sciences Sequencing Core for technical assistance with bulk RNA sequencing on the NextSeq 2000. We acknowledge Dr. Krista Gibbs, Heather Seiple, and Paige Prawucki in the Magee-Womens Research Institute animal facility for technical assistance in the care of mice. We also acknowledge the University of Pittsburgh Center for Research Computing through the resources provided. This work used the HTC cluster, supported by NIH award number S10OD028483. Lastly, we extend gratitude to Drs. William Walker, Christina Megli, Sandra Cascio, Kyle Orwig, Ronald Buckanovich, Judith Yanowitz, Mellissa Mann, Stephanie Correa, Tracy Bale, Jennifer Chan, Kristen Montgomery, Hannah Zierden, and Patrick Hanlin for their critical insights into this work.

## Author contributions

Conceptualization: SKM, JPG, AJZ, TWH, MAB-E, EJ

Methodology: SKM, JPG, AR, EW, NM, AK, YS, SJM, GEK, JDD, RJB, CCC, SLG, MG-L, KEM, TWH, MAB-E, EJ

Investigation: SKM, JPG, AR, EW, YS, SJM, JDD, RJB, CCC, GEK, KEM

Visualization: SKM, EJ Funding acquisition: EJ

Project administration: SKM, JPG, EJ Supervision: SKM, JPG, EJ

Writing – original draft: SKM, EJ

Writing – review & editing: SKM, JPG, AR, EW, NM, AK, YS, SJM, GEK, JDD, RJB, CCC, SLG, MG-L, KEM, AJZ, TWH, MAB-E, EJ.

## Data and materials availability

All data are available in the main text or the supplementary materials.

## Materials and Methods

### Mice

All experiments were approved by the University of Pittsburgh and Magee-Womens Research Institute Institutional Animal Care and Use Committee and performed in accordance with the National Institutes of Health Animal Care and Use Guidelines. Murine-pathogen-free C57Bl/6N female and male mice were purchased from Taconic Biosciences (C57Bl/6NTac). All mice were maintained on a 12-hour light/dark cycle (lights on 0600 EST; lights off 1800 EST). Room temperature was maintained between 75-77o F and 40-60% humidity. An Onset HOBO MX2202 Wireless Temperature/Light Data Logger (HOBO Data Loggers, Wilmington, NC) was used to confirm light-dark photoperiod stability. Unless otherwise specified, ad libitum access was provided to water and a grain-based rodent chow diet (NIH-31M Rodent Diet, Envigo; 20.940% protein, 65.324% carbohydrate, 13.736% fat) or a high-fat low-fiber diet (Research Diets D12492; 20.0% protein, 20.0% carbohydrate, 60.0% fat). Germ-free C57Bl/6NTac female and male mice were purchased from Taconic Biosciences and housed in the University of Pittsburgh Gnotobiotic Facility, with ad libitum access to water and the same grain-based rodent chow diet (NIH-31M Rodent Diet, Envigo; 20.940% protein, 65.324% carbohydrate, 13.736% fat).

### Confirmation of germ-free status

The University of Pittsburgh Gnotobiotic Facility conducts quarterly screening for bacterial contamination in germ-free mice. Screening for the detection of bacterial contamination of mouse feces by aerobic and anaerobic bacteria includes bacterial culture and qPCR assays. Screens were negative on all assays from isolators within which germ-free mice for this study were housed prior to the initiation of experiments and following the conclusion of the experiments. This confirmed the germ-free status of mice used in these studies.

### Germ-free mice microbial colonization

Two independent cohorts of time-mated pregnant germ-free dams were mated in four separate germ-free isolators during pregnancy. Each germfree isolator was randomly assigned to one of three groups: 1) pregnant dams that were conventionalized the afternoon prior to delivering offspring (designated as Ex-GF P0); 2) lactating dams that were colonized on day 12 post-delivery (designated as Ex-GF P12). For the microbial gavage preparation, ileal and cecal luminal contents were collected from 4 murine-pathogen-free C57Bl/6N Tac female mice at 8 weeks of age. Contents were weighed and transferred to a sterile tube with Rosshart Freezing Medium (RFM) and homogenized. Final gavage aliquot preparations were made at a 1:30 dilution (material: RFM), and this was used for microbiota associations. Female mice received one 200ul oral gavage of this preparation. Fecal pellets were collected from colonized mice at two-time points: 1) one day prior to gavage and 2) seven days following gavage.

### Short-chain fatty acid (SCFA) reconstitution

SCFA reconstitution experiments were conducted in one germ-free isolator. Time-mated germ-free dams were allowed to deliver and 12 days post-delivery, cages were randomized to receive drinking water supplemented with either a SCFA mix (67.5 mM sodium acetate, 40 mM sodium butyrate, 25,9 mM sodium propionate) or pH and salt-balanced vehicle control, following the protocol published in (*1*). The supplemented water was prepared fresh once a week, and the drinking water was changed weekly. Offspring remained on respective water treatment groups after weaning and until postnatal day 70 tissue collections.

### Tissue processing and Histological staining

Mouse ovaries were fixed in 4% paraformaldehyde (Thermo Scientific) at 4°C overnight. The tissue was then washed several times in calcium and magnesium-free PBS (Lonza). Washed tissues were paraffin-embedded, and 5µm sections were cut through the entire ovary with a microtome. Sections were mounted onto Superfrost-plus slides. Slides were stained with Hematoxylin-eosin or Periodic Acid Schiff (PAS) with the Leica BondMax Immunostaining System (Leica Biosystems). All tissue processing was performed by the Magee-Womens Research Institute histology and microimaging core. Masson’s Trichome staining was performed per manufacturer instructions using the Masson’s Trichome Stain Kit (Polysciences cat #25088).

### Microscopy

Image analysis was performed using Nikon Elements AR software on the Nikon 90i fully motorized imaging platform. Post-imaging analysis was performed using ImageJ V.2.14 software.

### Quantification of ovarian follicles

Follicle counts began from the first section, with ovarian tissue on every 10th section of serially sectioned tissue for the entire ovary (*2, 3*). Primordial follicles were classified as oocytes with a single layer of squamous granulosa cells. Primary follicles contained an oocyte surrounded by cuboidal granulosa cells. Oocytes partially surrounded by squamous and cuboidal granulosa cells were also counted as primary follicles. Secondary follicles contain an oocyte with more than one layer of granulosa cells. Antral follicles contained oocytes surrounded by multiple layers of granulosa and theca cells and a fluid-filled antrum. Corpus luteum contained granulosa lutein cells with the characteristic appearance of steroid-producing cells with pale cytoplasm and smaller, more deeply stained theca lutein cells. Atretic follicles were characterized by deeply stained zona pellucida detached from the granulosa cells with or without a fragmented oocyte and somatic cells with or without pyknotic nuclei. PAS staining confirmed the expression of zona pellucida protein on atretic follicles.

To prevent double counting, only follicles with an oocyte nucleus were counted. The number of follicles was determined by applying both the sampling frequency (10) and a correction factor to obtain a whole ovary and then a whole animal (× 2) value as previously described (*2–6*). The correction factor was determined as follows: (section thickness (5µm) ÷ (section thickness (5µm) + average follicle nucleus diameter. Oocyte nuclear diameters were measured in at least 10 representative follicles from each animal using Image J V.2.14 software.

### Superovulation and in vitro fertilization

Female mice from the CON, Hf-Lf, Early and Late diet intervention groups were injected interperitoneally (IP) with 15 IU pregnant mare serum gonadotropin (PMSG) (ProSpec #hor-272) at 5:30 PM on Day 1. On Day 3, forty-eight hours after PMSG, the mice received an IP injection of 15 IU of human chorionic gonadotropin (hCG) (Sigma #CG5). On Day 4, 9:00 AM, oocytes were harvested and pooled from the animals in each group (n=3-5/group). *In vitro* fertilization was performed using CARD Medium (Cosmo Bio USA #KYD-003-EX), and oocyte maturation and embryo developmental competence was evaluated daily on Days 5-7.

### Glucose tolerance test

Food was removed at 0800 EST and mice were fasted for 6 h. At 1400 EST, mice were injected intraperitoneally with 1 mg/kg BW of 0.3 g/mL glucose in saline. Glucose readings were collected by tail blood at 0-, 1530-, 60, and 120-minute time points using the Contour Next Blood Glucose Monitoring System (Bayer Co, Germany).

### 16S rRNA marker gene sequencing and data analysis

Genomic DNA was extracted from 50 mg of cecal luminal contents using the MagAttract PowerMicrobiome DNA/RNA Kit (Qiagen), with beadbeating on a TissueLyser II (Qiagen) according to the manufacturer’s instructions. 16S libraries were generated using a two-step PCR protocol. Amplicon PCR was performed as follows for amplification of the 16s rRNA V3-V4 region from cecal luminal contents: initial denaturation at 95°C for 3 minutes, followed by 25-cyles 95°C for 30 seconds, 55°C for 30 seconds, 72°C for 30 seconds, and a final extension at 72°C for 5 minutes. Resultant 16S V3-V4 amplicons were then purified using AMPure XP beads at a 0.8 ratio of beads to amplicon volume. Illumina Nextera XT v2 Index Primer 1 (N7xx) and Nextera XT v2 Index Primer 2 (S5xx) were index primers. Index PCR was performed as follows for amplification of the 16s rRNA V3-V4 region from cecal luminal contents: initial denaturation at 95°C for 3 minutes, followed by 8-cyles 95°C for 30 seconds, 55°C for 30 seconds, 72°C for 30 seconds, and a final extension at 72°C for 5 minutes. Results indexed libraries were cleaned up using AMPure XP beads at a 0.8 ratio of beads to the indexed library. The concentration of indexed libraries was quantified using a Qubit 4 fluorimeter, and library fragment size was quantified using an Agilent TapeStation 4200 with D5000 ScreenTapes. Libraries were normalized pooled, and a paired-end sequencing of pooled libraries was done on an Illumina iSeq 100 System using 2×150bp run geometry.

The sequences were demultiplexed on the BaseSpace Sequence Hub using the bcl2fastq2 conversion software (version 2.2.0.) and analyzed using the QIIME 2 (version 2022.2) microbiome bioinformatics platform (*7*). Quality control on the resulting demultiplexed forward fastq files was performed using DADA22 denoise-single function trimming 33bp of the primer sequence (*8*). A Naive Bayes feature classifier was trained using SILVA reference sequences (*9*) with the q2-feature-classifier for taxonomic analysis. The average count per sample was 27,611, with the maximum count per sample at 42,407 and the minimum count at 10,939. Statistical and metaanalysis of the data was conducted using MicrobiomeAnalyst (*10*). Data filtering was set to include features where 20% of its values contain a minimum of four counts. In addition, features that exhibit low variance across treatment conditions are unlikely to be associated with treatment conditions. Therefore, the variance was measured by the interquartile range and removed at 10%. Data were normalized by using center log ratio to account for compositional nature of these data. Taxa identified as cyanobacteria or ‘unclassified’ to the phylum level were removed.

### Whole shotgun metagenomics and data analysis

Genomic DNA extracted for 16S rRNA marker gene sequencing was utilized. Shotgun metagenomic libraries were prepared using the Illumina DNA pre kits according to the manufacturer’s protocol. The quality and quantity of the libraries were assessed using Qubit fluorometric quantitation and Agilent TapeStation 4200 with D5000 ScreenTapes. Sequencing was performed on an Illumina NextSeq 2000 platform, using 2 × 300 bp paired-end sequencing. A target sequencing depth of 40 million reads per sample was aimed for to ensure adequate coverage of the metagenomes. Raw sequencing data were processed using the WGSA2 (Whole Genome Shotgun Analysis 2) pipeline available on the Nephele platform (https://nephele.niaid.nih.gov/pipeline_details/wgsa/) (*11*). The pipeline was executed with default parameters unless otherwise specified. For quality control and preprocessing, raw reads were trimmed and filtered for quality using Trimmomatic (*12*). For host DNA removal, sequences matching the mouse genome were removed using Bowtie2 (*13*). Filtered reads were classified using Kraken2 (*14*), and protein-coding sequences were predicted and annotated using Prodigal (*15*) and compared against the KEGG database (*16, 17*).

### Quantification of 3NP-Short Chain Fatty Acids

Cecal luminal contents and ovarian tissue samples were homogenized with 50% aqueous acetonitrile at a ratio of 1:15 vol:wt. Deuterated internal standards (5 µg/mL): formate-d2, acetate-d4, butyrate-d5, proprionate-d6, valerate-d2 and hexanoate-d4 (CDN Isotopes, Quebec, Canada) were added. Samples were homogenized using a FastPrep-24 system (MP-Bio), with Matrix D at 60 Hz for 30 seconds, before being cleared of protein by centrifugation at 16,000xg. Plasma samples were cleared of protein using 4x volumes ice-cold 1:1 MeOH:EtOH with vortexing, followed by centrifugation at 16,000xg. Cleared supernatant (60 µL) was collected and derivatized using 3-nitrophenylhydrazine (3-NP). Each sample was mixed with 20 µL of 200 mM 3-nitrophenylhydrazine in 50% aqueous acetonitrile and 20 µL of 120 mM N-(3-dimethylaminopropyl)-N’-ethyl carbodiimide 6% pyridine solution in 50% aqueous acetonitrile. The mixture reacted at 60°C for 40 minutes, and the reaction was stopped with 0.45 mL of 50% acetonitrile. Absolute quantitation of SCFAs was performed by preparing 9-point calibration curves for formate (1.102 fmoloµL to 0.241 nmol/µL) acetate (0.8 fmol/µL to 0.185 nmol/µL), propionate (0.6 fmol/µL to 0.150 nmol/µL), and butyrate (0.5fmol/µL to 0.126 nmol/µL) and were derivatized as described above. Derivatized samples were injected (5 µL) via a Thermo Vanquish UHPLC and separated over a reversed-phase Phenomenex Kinetex 150 mm x 2.1 mm 1.7 µM particle C18 maintained at 55°C. For the 20-minute LC gradient, the mobile phase consisted of solvent A (water/0.1% FA) and solvent B (acetonitrile/0.1% FA). The gradient started at 15% B for 2 minutes, increased to 60% B over 10 minutes, and then to 100% B over 1 minute. The gradient was held at 100% B for 3 minutes, before equilibration at 15% B for 4 minutes. The Exploris 240 hybrid mass spectrometer was operated in positive ion mode, scanning in ddMS2 mode (2 µscans) from 75 to 1000 m/z at 120,000 resolution with an AGC target of 2e5 for full scan, 2e4 for MS2 scans using HCD fragmentation at stepped 15,35,50 collision energies. The source ionization setting was 3.0 kV spray voltage for positive mode. Source gas parameters were 45 sheath gas, 12 auxiliary gas at 320°C, and 3 sweep gas. Calibration was performed before analysis using the PierceTM FlexMix Ion Calibration Solutions (Thermo Fisher Scientific). Integrated peak areas were then extracted manually using Quan Browser (Thermo Fisher Xcalibur ver. 2.7) for both SCFA and organic acids (*18*). SCFAs are reported as the area ratio of SCFA to the internal standard before conversion to absolute concentration. Organic acid peak areas were normalized to the butyrate-d5 internal standard and reported as Relative Abundance.

### Whole-tissue RNA sequencing and data analysis

Frozen tissue samples were homogenized in QIAzol Reagent (Qiagen) using a MiltenyiBiotec GentleMACS Octo Dissociator for 30s. RNA was isolated with Qiagen miRNeasy Mini Kits according to the manufacturer’s instructions. RNA integrity was quantified on an Agilent TapeStation 4200 using TapeStation RNA ScreenTapes. All samples had an RIN score above 8. Sequencing libraries were prepared using Illumina Stranded mRNA prep, Ligation kits with IDT for Illumina RNA UD Indexes Set A, and Ligation index adapters. The concentration of indexed libraries was quantified using Qubit, and library fragment size was quantified using an Agilent TapeStation 4200 with D5000 ScreenTapes. Sequencing was performed on an Illumina NextSeq 2000 using P3 flow cells and 2×100 paired end-run geometry at the Health Sciences Sequencing Core at Children’s Hospital of Pittsburgh. Sequencing was repeated twice on the same library pool to achieve sufficient resolution and minimize batch effects, yielding an average of 30 – 60 million reads per sample.

### Whole-tissue RNA sequencing and data analysis

Frozen tissue samples were homogenized in QIAzol Reagent (Qiagen) using a MiltenyiBiotec GentleMACS Octo Dissociator for 30s. RNA was isolated with Qiagen miRNeasy Mini Kits according to the manufacturer’s instructions. RNA integrity was quantified on an Agilent TapeStation 4200 using TapeStation RNA ScreenTapes. All samples had an RIN score above 8. Sequencing libraries were prepared using Illumina Stranded mRNA prep, Ligation kits with IDT for Illumina RNA UD Indexes Set A, and Ligation index adapters. The concentration of indexed libraries was quantified using Qubit, and library fragment size was quantified using an Agilent TapeStation 4200 with D5000 ScreenTapes. Sequencing was performed on an Illumina NextSeq 2000 using P3 flow cells and 2×100 paired end-run geometry at the Health Sciences Sequencing Core at Children’s Hospital of Pittsburgh. Sequencing was repeated twice on the same library pool to achieve sufficient resolution and minimize batch effects, yielding an average of 30 – 60 million reads per sample.

Concatenated FASTQ files generated from Illumina were used as input to kallisto (*19*), a program that pseudo-aligns high-throughput sequencing reads to the Mus musculus reference transcriptome (version 38) and quantifies transcript expression. We used 60 bootstrap samples to ensure accurate transcript quantification. Gene isoforms were collapsed to gene symbols using the Bioconductor package tximport (version 3.4). Genes were filtered to counts per million >1 in at least three samples. The filtered gene list was normalized using a trimmed mean of M-values in edgeR (*20*), modeled mean-variance trends and fit linear model to RNAseq data using voom (*21*), and differential gene expression analysis was conducted using linear modeling using limma (*22*). Volcano plots and heatmaps were visualized using ggplots (*23*) and heatmaply (*24*).

### Time-series analysis of whole-tissue RNA-seq data

To model genes that vary in their expression across postnatal stages, we used the algorithm splineTimeR (*25, 26*). As input for splineTimeR (v1.1.0), we used normalized log2 transformed counts. This tool fits natural cubic spline curves to time-course data and applies empirical Bayes moderate F-statistics on the coefficients of the spline regression model between two groups for detecting differentially expressed genes over time. We used SplineDiffExprs() with intercept=TRUE and df=4 for detecting microbiota-dependent genes. P-values were adjusted for multiple testing using the Benjamini-Hochberg (BH) procedure, with an adjusted P < 0.25 indicating significance.

### TaqMan assays of the arcuate nucleus of the hypothalamus

TaqMan gene expression analysis of the adult arcuate nucleus of the hypothalamus (Arc). Frozen brains from P70 female mice were cryosectioned at −20 °C. Using a hollow 1.0 mm needle, the Arc was removed according to the mouse brain atlas. Arc micropunches were immediately dispensed into 500 µl of Trizol and stored at −80 °C until processing. Messenger RNA was purified RNeasy Mini kit (Qiagen).

### Quantification and statistical analysis

Statistical information, including sample size, mean, and statistical significance values, are shown in the text or the figure legends. A variety of statistical analyses were applied, each one specifically appropriate for the data and hypothesis as depicted in corresponding figures, using GraphPad Prism 9.3.1 or R version 4.1.0 (R Core Team, 2021). Processing of RNAseq data was conducted using standardized and published protocols. GraphPad Prism and Adobe Illustrator were used for generating figures. No custom script was used to analyze RNA sequencing data.

## Supplementary Materials for

**Fig. S1.**
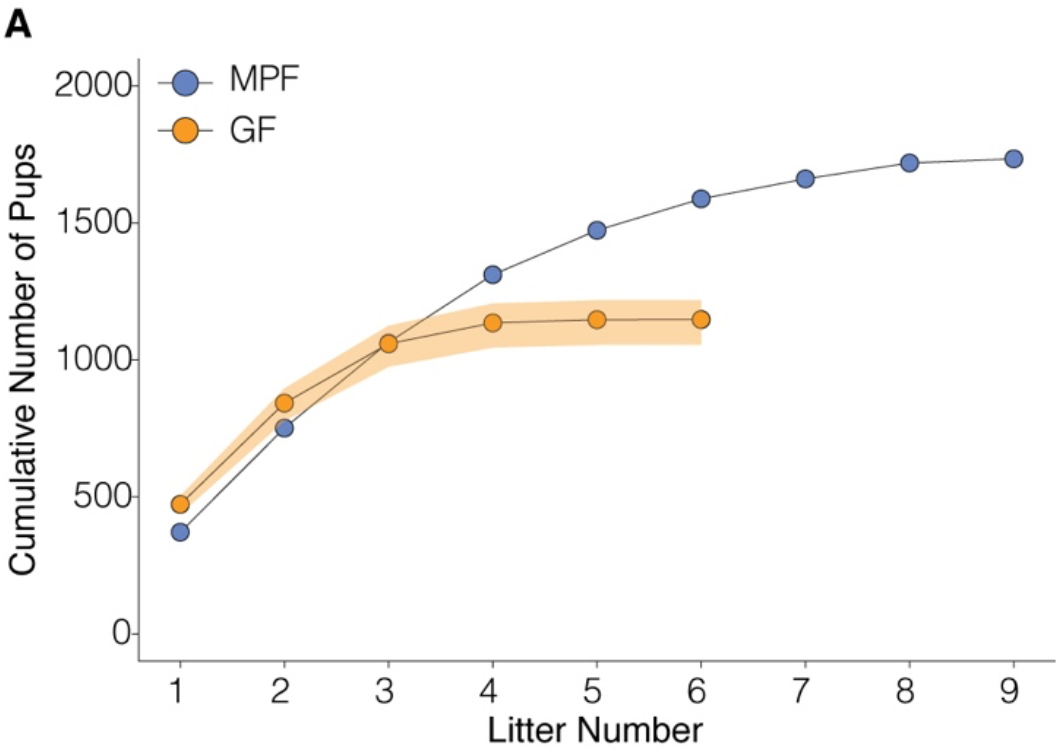
Undersampling analysis of cumulative pup production in MPF and GF mice across sequential litters. (**A**) Undersampling analysis (1000 permutations) of cumulative number of pups produced by MPF (blue) and GF (orange) female mice over successive litters. Each point represents the mean cumulative pup count at a given litter number. Shaded areas indicate 95% confidence intervals. MPF mice show continued reproductive output beyond the 6th litter, while GF mice reproduction appears to plateau after the 5th litter. n = 58 MPF and 89 GF females, total of 493 litters. Data points represent means. Shaded areas represent 95% confidence intervals. MPF, murine pathogen free; GF, germ-free. See Figure 1 for additional reproductive output data.

**Fig. S2.**
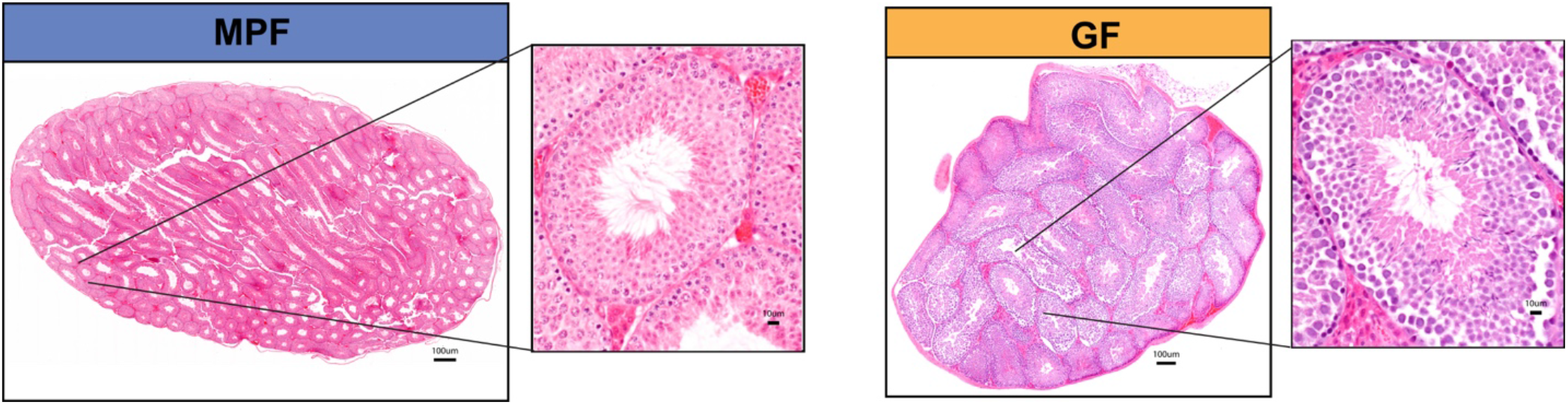
Comparable testicular histology in 10-week-old MPF and GF male mice. Representative testicular histology from 10-week-old MPF (left) and GF (right) male mice. H&E staining, scale bars: 100 μm. Insets show higher magnification of seminiferous tubules (scale bars: 10 μm). Both MPF and GF testes display normal spermatogenesis, including the presence of spermatids in the lumen of seminiferous tubules.

**Fig. S3.**
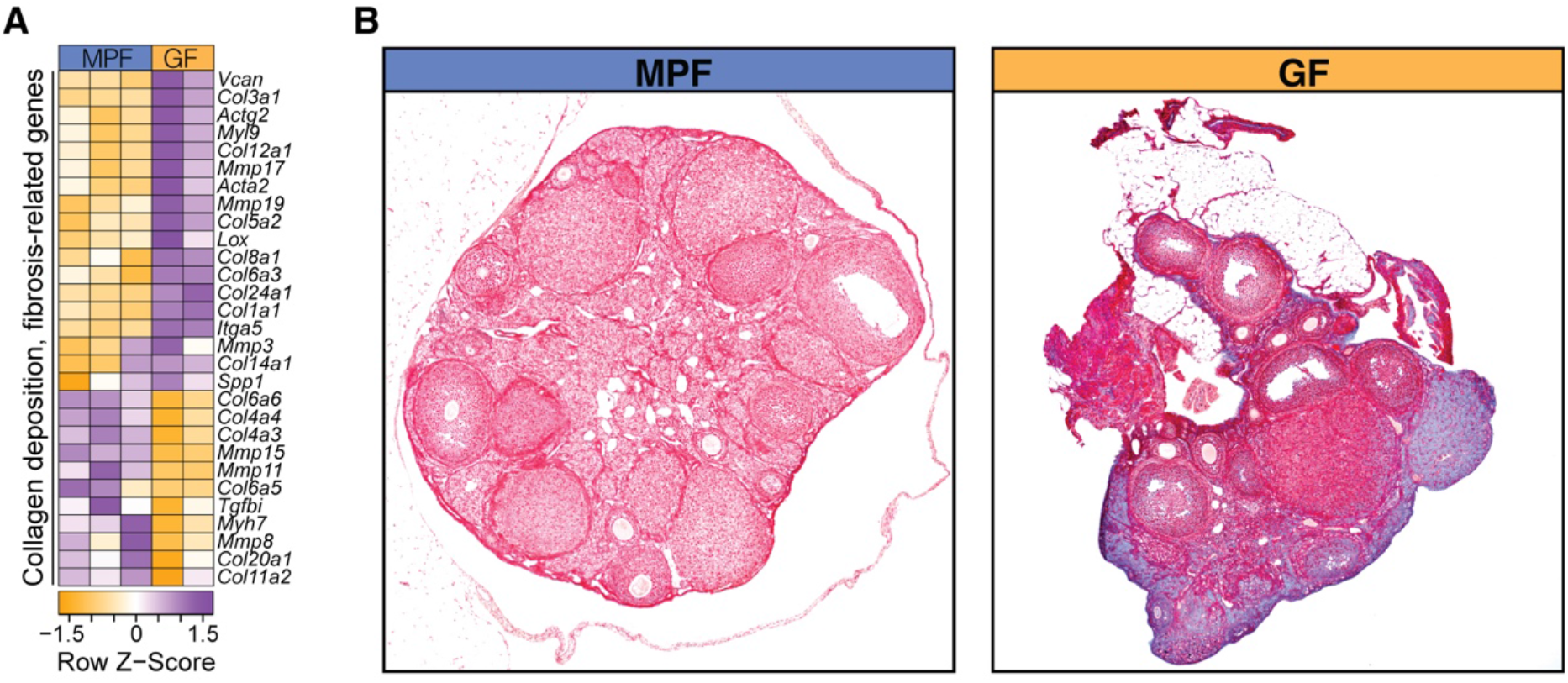
Gut microbiota regulates ovarian extracellular matrix composition. (**A**) Heatmap of collagen deposition and fibrosis-related gene expression in MPF and GF ovaries at 10 weeks (FDR < 0.25). Data displayed as row Z-scores. GF ovaries show higher expression of several collagen and ECM-related genes compared to MPF. (**B**) Representative Masson’s trichrome staining of ovarian sections from MPF (left) and GF (right) mice. Blue staining indicates collagen deposition. GF ovary displays increased collagen deposition and fibrotic changes compared to MPF. n = 3 MPF and 2 GF samples for RNA-seq. n = 3-5 mice per group for histology. MPF, murine pathogen free; GF, germ-free; ECM, extracellular matrix. See Figure 2 for related ovarian histology and gene expression data.

**Fig. S4.**
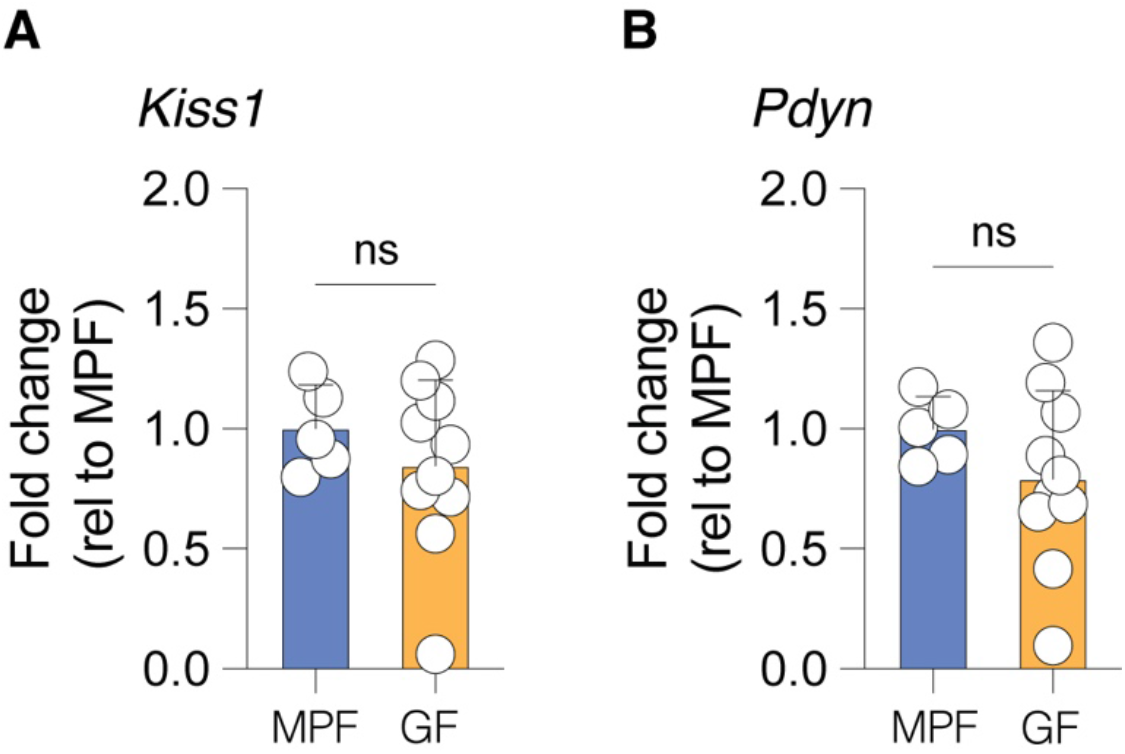
Comparable *Kiss1* and *Pdyn* expression in the arcuate nucleus of the hypothalamus of MPF and GF female mice at P70. **(A)**Fold change in *Kiss1* mRNA transcript levels in the arcuate nucleus of MPF and GF mice. **(B)**Fold change in *Pdyn* mRNA transcript levels in the arcuate nucleus of MPF and GF mice. n = 5-10 mice per group. Data: mean ± SD, analyzed using unpaired t-test. ns, not significant. MPF, murine pathogen free; GF, germ-free.

**Fig. S5.**
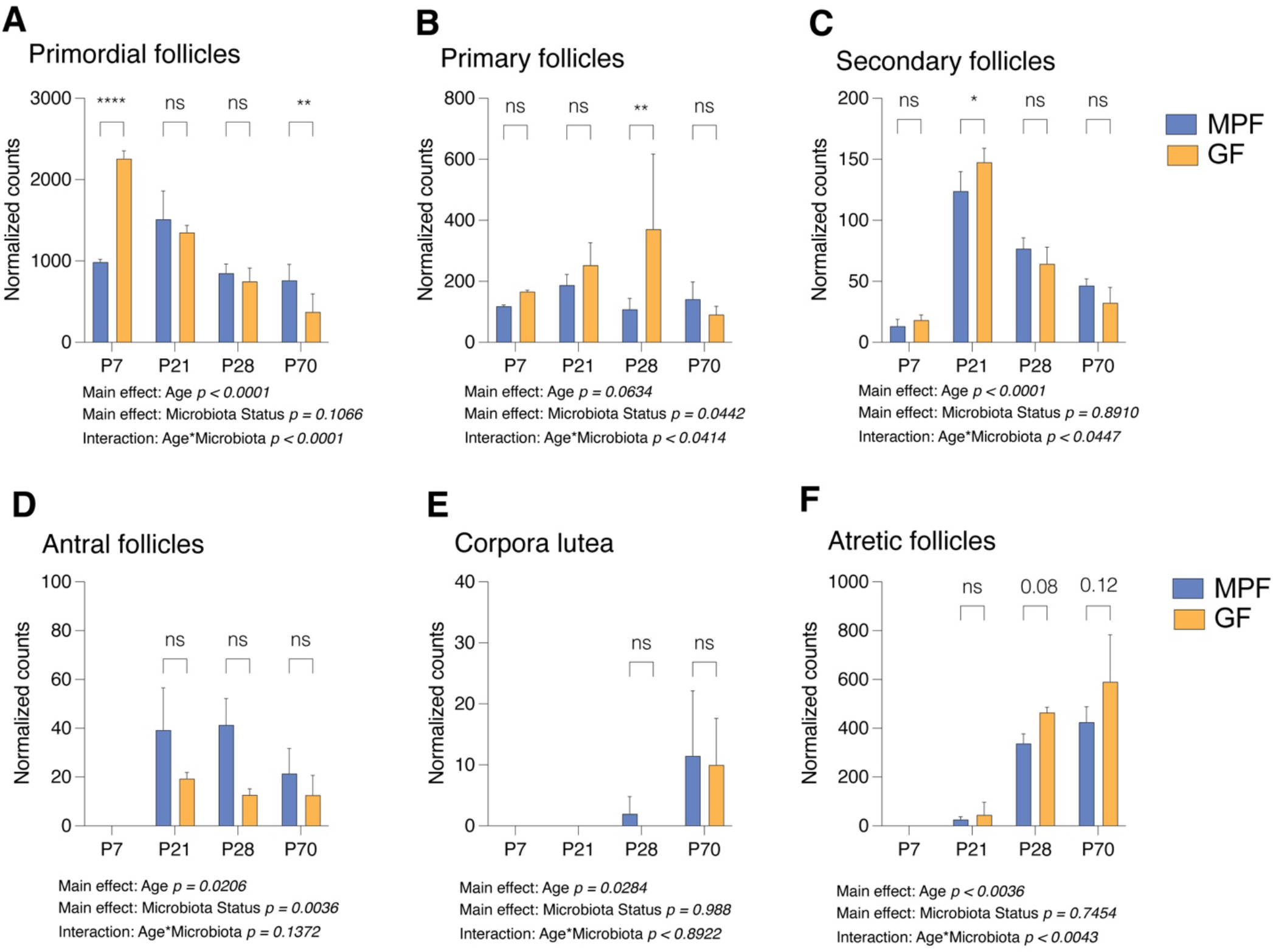
Gut microbiota influences ovarian follicle dynamics during postnatal development. (**A-F**) Quantification of ovarian follicles in MPF and GF mice across postnatal development (P7-P70): (**A)**Primordial follicles. (**B)**Primary follicles. (**C)**Secondary follicles. (**D)**Antral follicles. (**E)**Corpora lutea. (**F)**Atretic follicles. n = 3-5 per group and timepoint. Data: mean ± SD, analyzed by two-way ANOVA followed by Fisher’s LSD test (A-D), analyzed by Poisson regression with Tukey’s test (E-F). Main effects of Age, Microbiota Status, and their interaction are reported for each follicle type. *p < 0.05, **p < 0.01, ****p < 0.0001, ns = not significant. MPF, murine pathogen free; GF, germ-free. See Figure 3 for related ovarian follicle dynamics data.

**Fig. S6.**
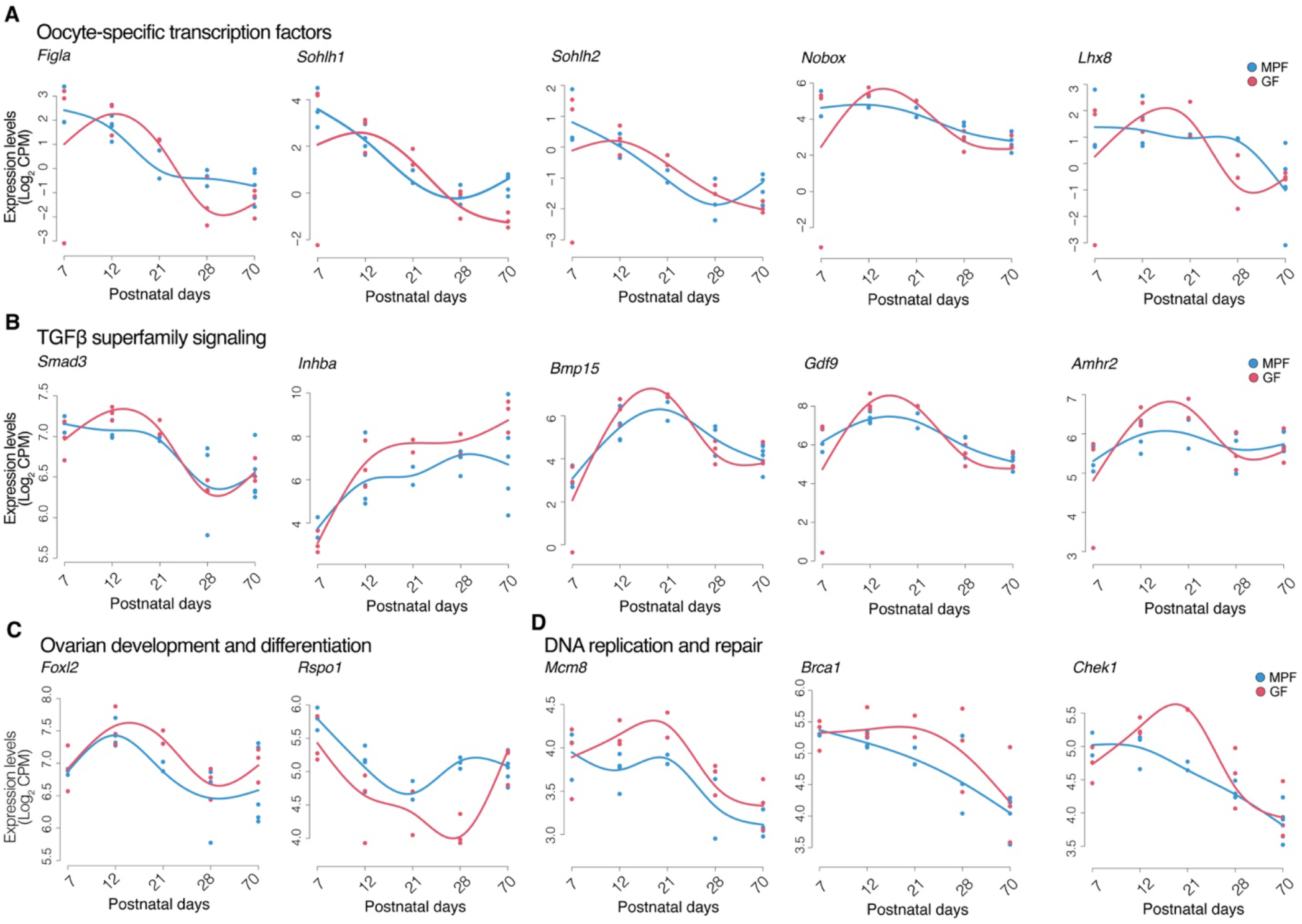
Gut microbiota modulates ovarian gene expression profiles during postnatal development. (**A-D**) RNA-seq analysis of ovarian gene expression in MPF (blue) and GF (red) mice at P7, P12, P21, P28, and P70: (**A)**Oocyte-specific transcription factors: *Figla, Sohlh1, Sohlh2, Nobox, Lhx8* (**B)**TGFβ superfamily signaling: Smad3, *Inhba, Bmp15, Gdf9, Amhr2* (**C)**Ovarian development and differentiation: *Foxl2, Rspo1* (**D)**DNA replication and repair: *Mcm8, Brca1, Chek1* n = 3-5 samples per group and timepoint. Data: filtered, normalized, log2 counts per million. Each dot represents an individual sample; lines show time-series mean expression trends. MPF, murine pathogen free; GF, germ-free. See Figure 3 for related ovarian gene expression data.

**Fig. S7.**
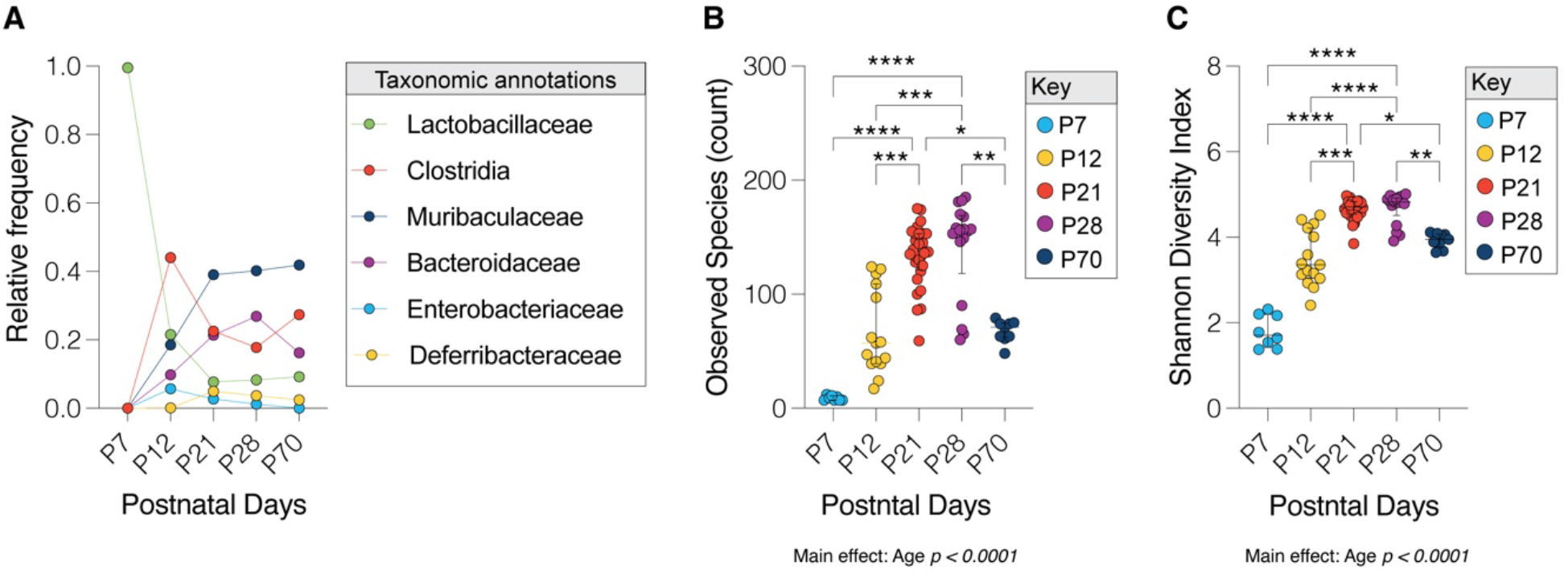
Cecal microbiota composition and diversity dynamics during postnatal development in MPF female mice. **(A)** Relative frequency of major microbial families in cecal lumen from P7 to P70. **(B)**Alpha diversity measured by observed species count (richness) across postnatal development. **(C)** Alpha diversity measured by Shannon diversity index across postnatal development. n = 8-28 mice per timepoint. Data in (B) and (C): median ± interquartile range, analyzed by Kruskal-Wallis test followed by Dunn’s test. *p < 0.05, **p < 0.01, ***p < 0.001, ****p < 0.0001. MPF, murine pathogen free. See Figure 3 for related microbiota analysis.

**Fig. S8.**
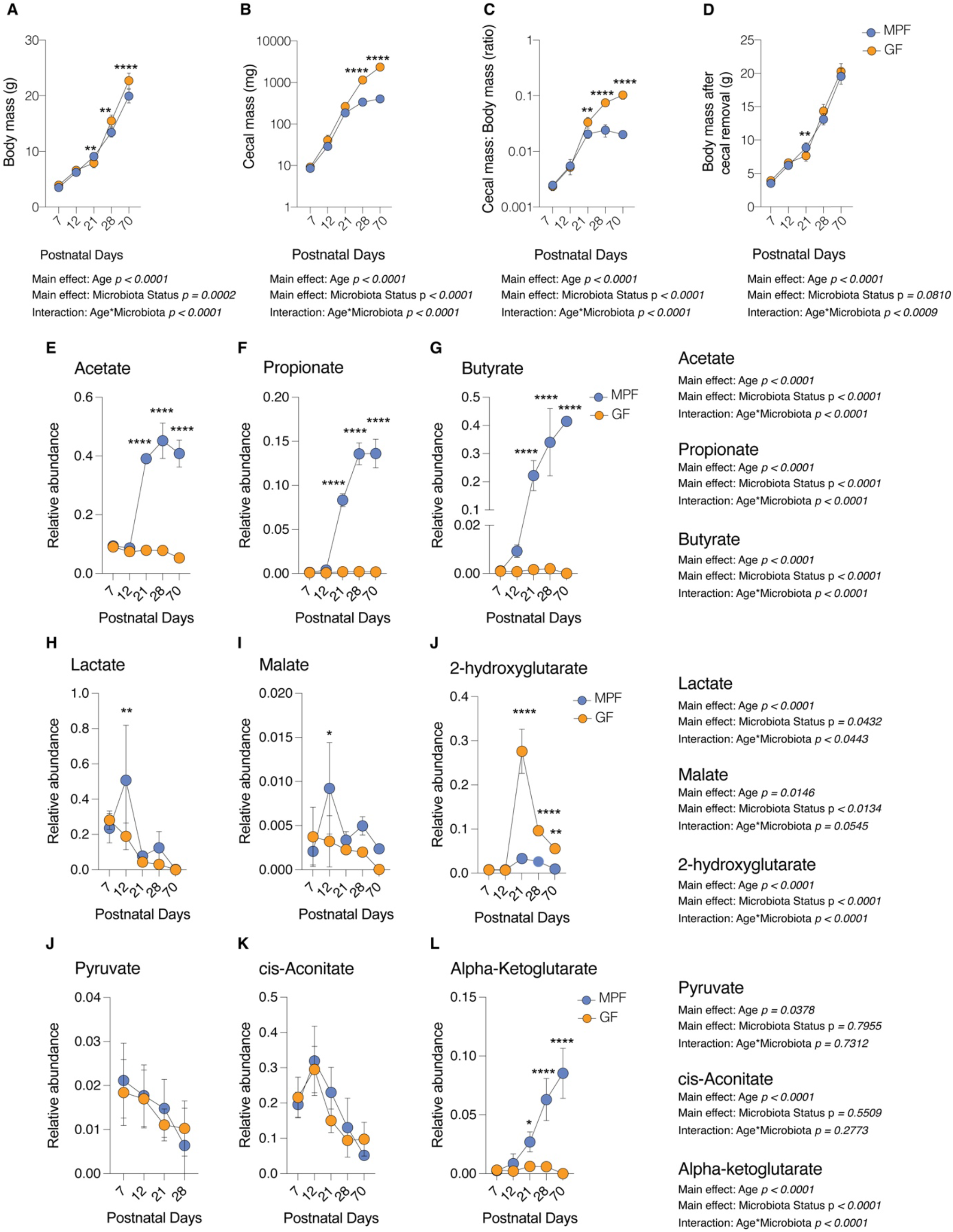
Gut microbiota influences metabolic parameters across postnatal development. (**A-D**) Whole body parameters of MPF and GF female mice: **(A)** Body mass **(B)** Cecal mass **(C)** Cecal mass to body mass ratio **(D)** Body mass after cecal removal (**E-G**) Short-chain fatty acids in cecal contents: **(E)** Acetate **(F)** Propionate **(G)** Butyrate (**H-M**) TCA cycle intermediates in cecal contents: **(H)** Lactate **(I)**Malate **(J)**2-hydroxyglutarate **(K)**Pyruvate **(L)**cis-Aconitate **(M)** Alpha-ketoglutarate n = 3-16 mice per group and timepoint (A-D); n = 3-5 mice per group and timepoint (E-M). Data: mean ± SD, analyzed by two-way ANOVA followed by Šidák’s test. Main effects of Age, Microbiota Status, and their interaction are reported for each comparison. *p < 0.05, **p < 0.01, ***p < 0.001, ****p < 0.0001. MPF, murine pathogen free; GF, germ-free. See Figure 3 for related metabolic data.

**Fig. S9.**
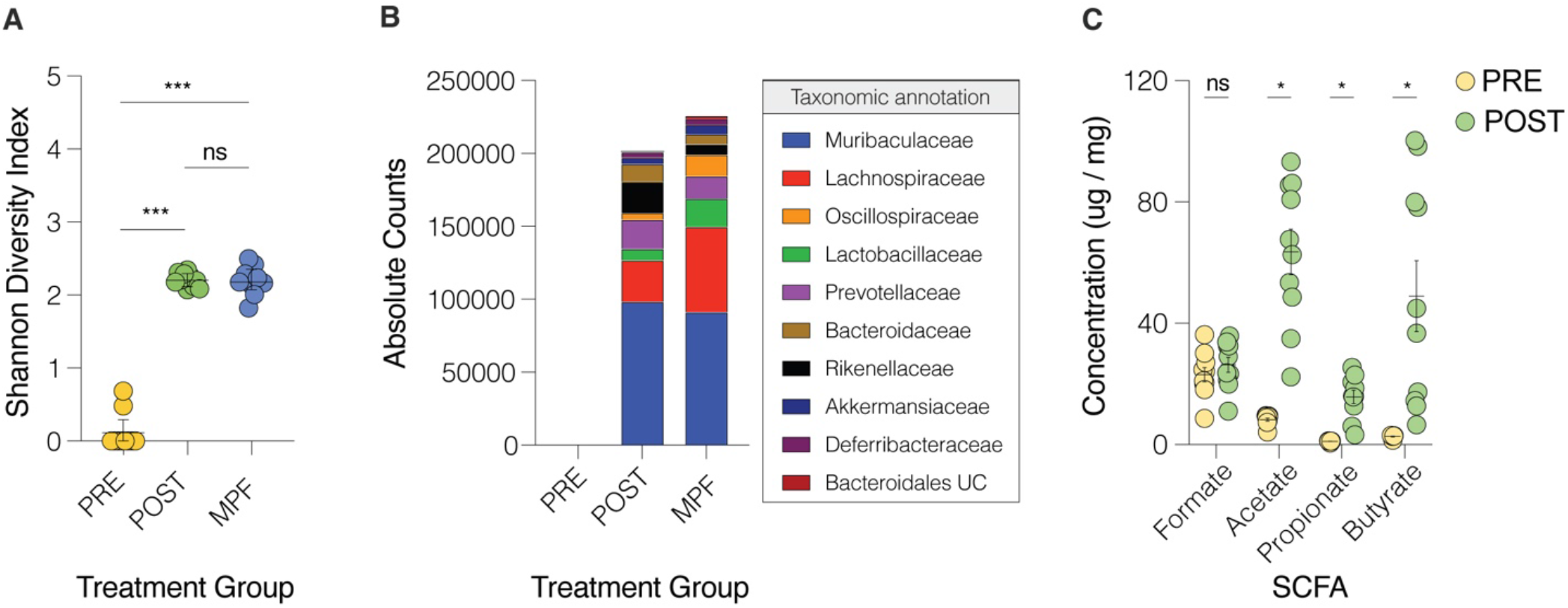
Successful microbiota engraftment in colonized GF dams. **(A)** Shannon Diversity Index of cecal microbiota in PRE-colonization, POST-colonization, and MPF female mice. **(B)** Absolute counts of major bacterial taxa in cecal microbiota of PRE-colonization, POST-colonization, and MPF female mice. **(C)** Concentrations of SCFAs in cecal contents of PRE-colonization and POST-colonization female mice. n = 8-10 mice per group. Data: mean ± SD. Statistics: (A) one-way ANOVA followed by Tukey’s test, (C) unpaired t-test. *p < 0.05, ***p < 0.001, ns = not significant. PRE, before colonization; POST, 7 days post-colonization; MPF, murine pathogen free; SCFA, short-chain fatty acids. See Figure 4 for related colonization data.

**Fig. S10.**
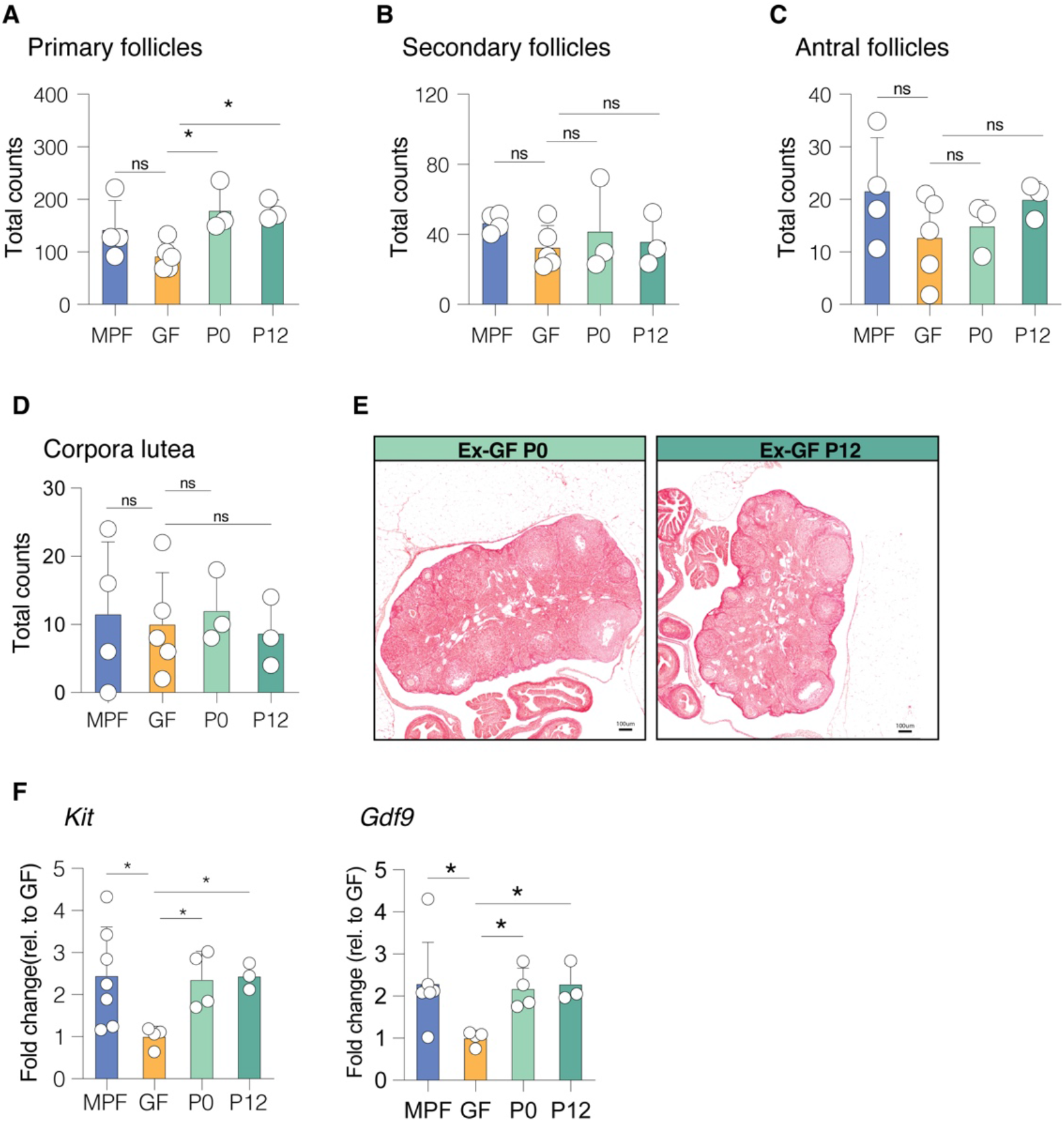
Early-life microbiota colonization mediates ovarian follicle dynamics. (**A-D**) Quantification of ovarian follicles in MPF, P0-colonized, P12-colonized, and GF female mice at P70: **(A)**Primary follicles **(B)** Secondary follicles **(C)** Antral follicles **(D)** Corpora lutea **(E)** Representative Masson’s Trichrome stained ovarian sections from P0-colonized and P12-colonized female mice at P70. **(F)** qPCR of ovarian tissue *Kit* and *Gdf9* mRNA transcript levels. n = 3-5 mice per group. Data: mean ± SD, analyzed by one-way ANOVA followed by Fisher’s LSD test (A-C), analyzed using Poisson regression with Tukey’s test (D). *p < 0.05, ns = not significant. MPF, murine pathogen free; P0, colonized at birth; P12, colonized at postnatal day 12; GF, germ-free. Scale bars: 100 μm. See Figure 4 for related ovarian follicle data and Figure S3 for GF ovarian histology.

**Fig. S11.**
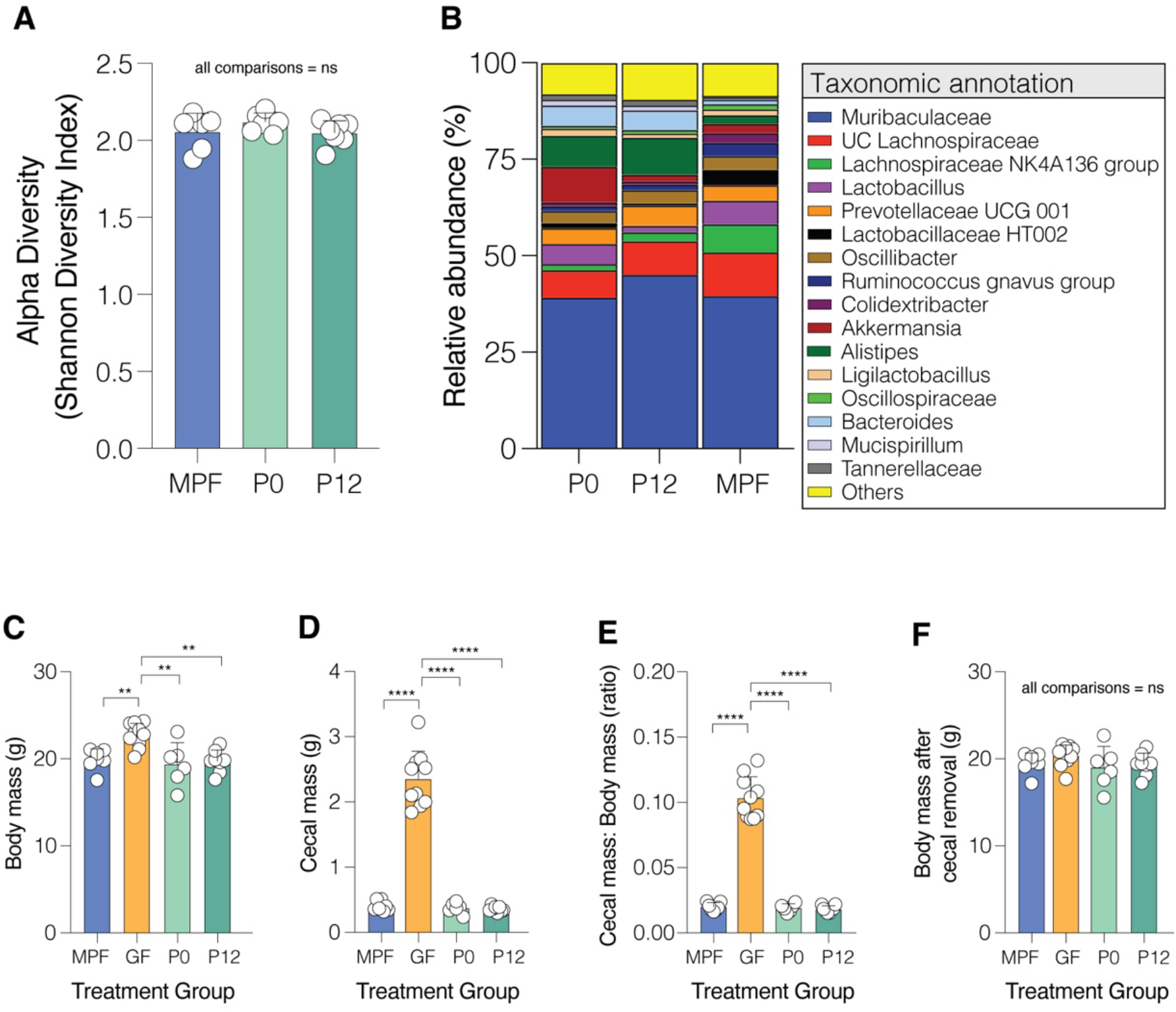
Early-life microbiota colonization normalizes cecal microbial composition and metabolic parameters. **(A)** Alpha Diversity (Shannon Diversity Index) of cecal microbiota in P0-colonized, P12-colonized, and MPF female mice at P70. **(B)**Relative abundance of major bacterial taxa in cecal microbiota of P0-colonized, P12-colonized, and MPF female mice at P70. (**C-F**) Metabolic parameters in MPF, GF, P0-colonized, and P12-colonized female mice at P70: **(C)**Body mass **(D)**Cecal mass **(E)**Cecal mass to body mass ratio **(F)** Body mass after cecal removal n = 6-8 mice per group. Data: mean ± SD. Statistics: (A) Kruskal-Wallis test, (C-F) One-way ANOVA followed by Tukey’s test. ns = not significant, **p < 0.01, ****p < 0.0001. MPF, murine pathogen free; GF, germ-free; P0, colonized at birth; P12, colonized at postnatal day 12. See Figure 4 for related colonization data.

**Fig. S12.**
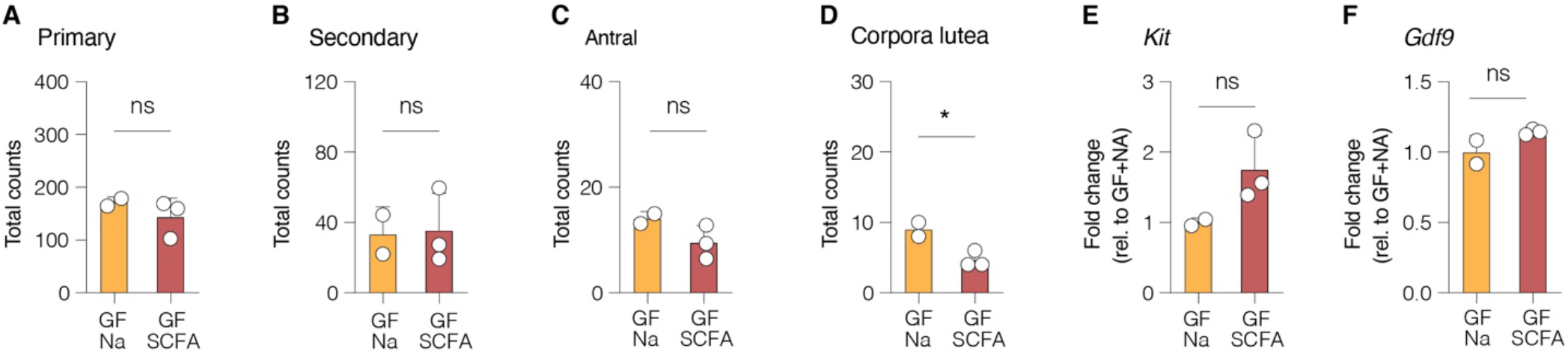
SCFA supplementation effects on ovarian follicles and ovary gene expression. (**A-D**) Quantification of ovarian follicles in GF+Na (Vehicle) and GF+SCFA female mice at P70: **(A)** Primary follicles **(B)** Secondary follicles **(C**) Antral follicles **(D)** Corpora lutea (**E**,**F**) qPCR of ovarian tissue *Amh, Nobox*, and *Gdf9* mRNA transcript levels. n = 2-3 mice per group. Na, pH and sodium matched Vehicle water; SCFA, short-chain fatty acids. See Figure 4 for related ovarian follicle counts and ovarian qPCR data.

**Fig. S13.**
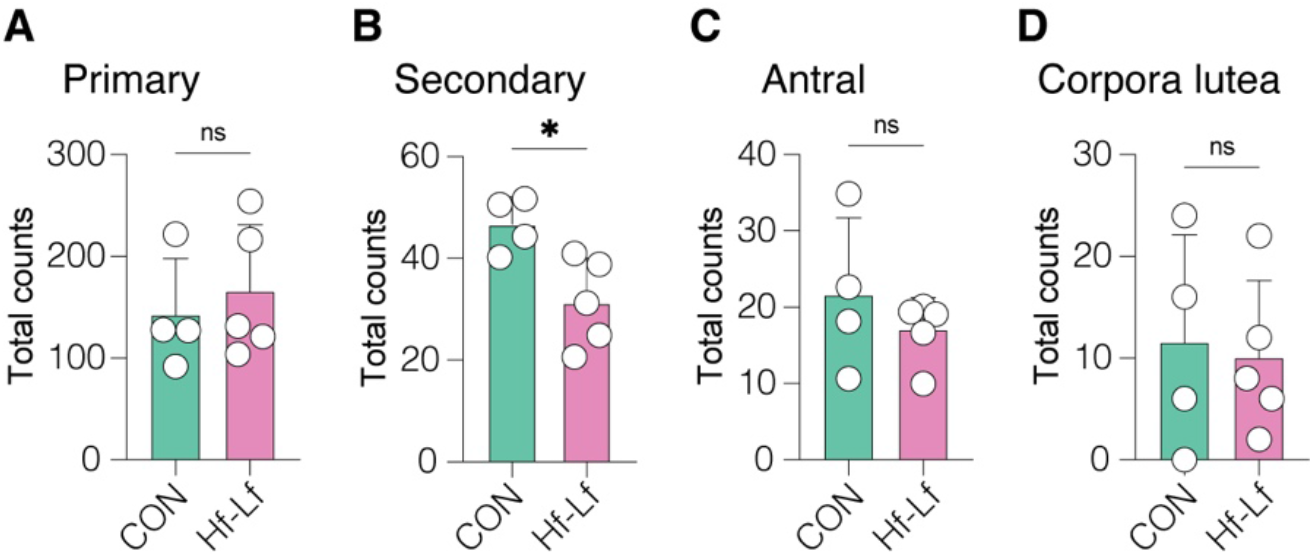
High-fat, low-fiber diet effects on ovarian follicles at P70 is primarily driven by effects on primordial and atretic follicle counts. (**A-D**) Quantification of ovarian follicles in control (CON) and high-fat, low-fiber diet-fed (Hf-Lf) female mice at P70: **(A)**Primary follicles **(B)**Secondary follicles **(C)**Antral follicles **(D)**Corpora lutea n = 4-5 mice per group. Data: mean ± SD, analyzed using unpaired t-test (A-C), analyzed using Poisson regression with Tukey’s test (D). *p < 0.05, ns = not significant. See Figure 5 for related ovarian follicle data.

**Fig. S14.**
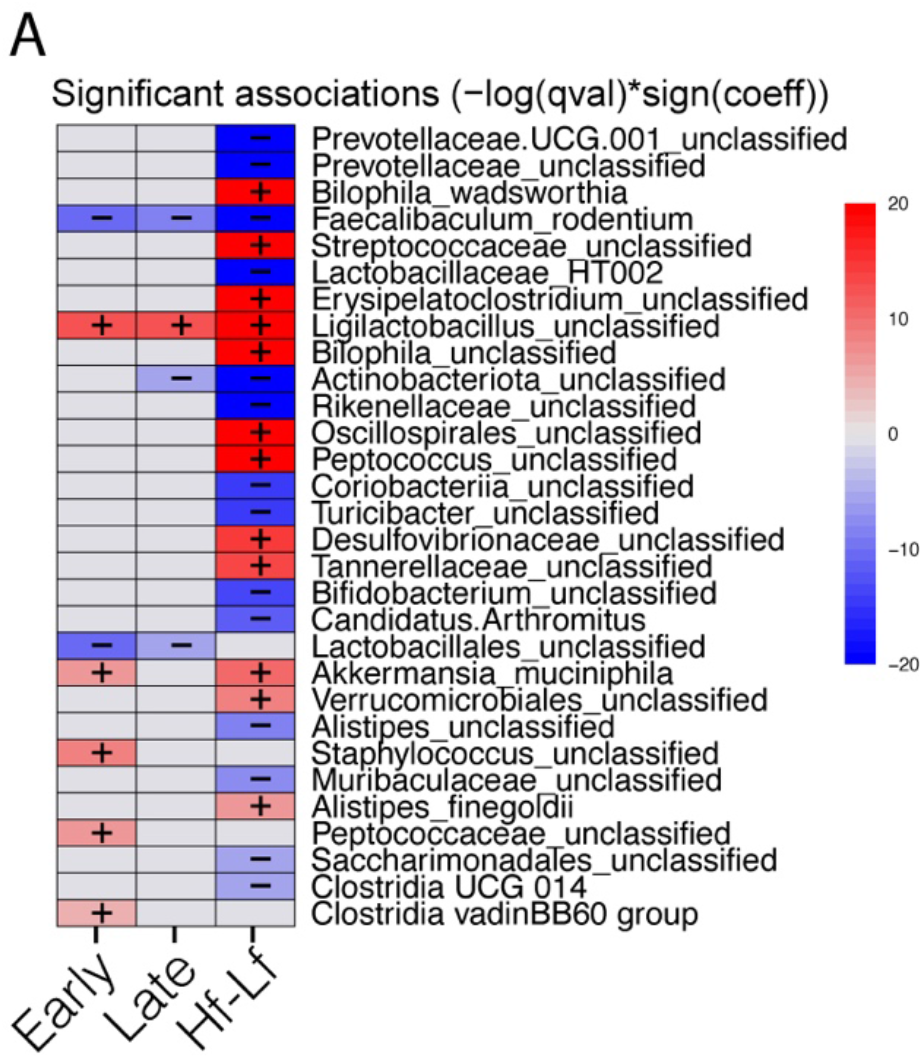
Differential abundance analysis of gut microbiota in response to dietary interventions. **(A)**Heatmap showing significant associations between bacterial taxa and diet intervention groups (Early, Late, and Hf-Lf) relative to the CON diet, using MaAsLin2 (Microbiome Multivariable Association with Linear Models 2) differential abundance analysis. Colors represent the strength and direction of associations compared to CON, with red indicating increased abundance and blue indicating decreased abundance. The intensity of the color corresponds to the magnitude of the association, calculated as -log(qval)*sign(coeff). Grey cells indicate non-significant differences from CON. Taxa names are listed on the right, while diet intervention groups are shown at the bottom. The color scale ranges from -20 (dark blue, strongly decreased) to 20 (dark red, strongly increased) relative to CON. See Figure 5 for related microbiota data.

